# Semisupervised definition of hippocampal ripples

**DOI:** 10.1101/2020.10.23.353102

**Authors:** Yusuke Watanabe, Mami Okada, Yuji Ikegaya

## Abstract

Hippocampal ripples are transient neuronal features observed in high-frequency oscillatory bands of local field potentials, and they occur primarily during periods of behavioral immobility and slow-wave sleep. Ripples have been defined based on mathematically engineered features, such as magnitudes, durations, and cycles per event. However, the “ripples” could vary from laboratory to laboratory because their definition is subject to human bias, including the arbitrary choice of parameters and thresholds. In addition, local field potentials are often influenced by myoelectric noise arising from animal movement, making it difficult to distinguish ripples from high-frequency noises. To overcome these problems, we extracted ripple candidates under few constraints and labeled them as binary or stochastic “true” or “false” ripples using Gaussian mixed model clustering and a deep convolutional neural network in a weakly supervised fashion. Our automatic method separated ripples and myoelectric noise and was able to detect ripples even when the animals were moving. Moreover, we confirmed that a convolutional neural network was able to detect ripples defined by our method. Leave-one-animal-out cross-validation estimated the area under the precision-recall curve for ripple detection to be 0.72. Finally, our model establishes an appropriate threshold for the ripple magnitude in the case of the conventional detection of ripples.

## Introduction

Ripples are one form of the local field potential (LFP) oscillations observed in the hippocampus. As the name “ripple” suggests, hippocampal ripples were first recognized based on their visually distinctive waveforms; they are characterized by 150–250 Hz components and durations of < 150 ms (O’Keefe, 1976). Ripples mainly occur during periods of awake immobility, consummatory behavior, and slow-wave sleep (Buzsáki et al., 1992), thereby contributing to memory consolidation (Girardeau et al., 2009; Ego-Stengel and Wilson, 2010) and relating to memory retrieval (Wu et al., 2017; Norman et al., 2019). In conventional strategies for the detection of ripples in LFPs, ripples are determined based on criteria such as the ripple band root mean square (RMS) amplitude, duration, cycles per event, and animal head speed (Ramirez-Villegas et al., 2015; Karlsson and Frank, 2009; Kay et al., 2016; Fernández-Ruiz et al., 2019; Shin et al., 2019; Hannah et al., 2019). In this process, analysts need to define ripples by arbitrarily setting the thresholds of these parameters. Therefore, the thresholds often vary among researchers, and “ripples” are inconsistent among laboratories, potentially leading to different conclusions.

Another problem is that the thresholds are usually predetermined and fixed in experiments, although the LFP waveforms may differ among animals, recording sites, and electrodes. Thus, in some laboratories, including those of our research group, when detected, ripples are manually screened by eye (Ramirez-Villegas et al., 2015; Norimoto et al., 2018). Eye inspection requires a great deal of skill and is subject to human bias. This task is also laborious and hinders upscaling to large data sets. Thus, overall, the current method of detecting ripples has problems related to objectivity, consistency, and reproducibility.

Deep convolutional neural networks (CNNs) are, in general, an appropriate tool for capturing the shapes or local features of objects. A CNN is a mathematical model inspired by the visual cortex system (Fukushima, 1980; Krizhevsky et al., 2012). CNNs have been studied since the ILSVRC2012 (ImageNet Large Scale Visual Recognition Challenge 2012), and the ability of these models to classify images is beyond that of humans (He et al., 2016). In the present study, we used a one-dimensional CNN to define ripples with reduced human bias. The one-dimensional CNN extracts local features from LFP signals in the time domain and learns to discriminate true and false ripples at a stochastic scale. Additionally, based on the defined ripples, we evaluated the ability of the CNN to detect a ripple event from a series of 400-ms periods of raw LFPs.

## Materials and Methods

### Animal Ethics

All animal experiments were performed following the University of Tokyo Animal Experiments Implementation Manual with the Animal Experiment Committee’s approval to minimize pain for experimental animals (approval number: P29-14). All mice were housed based on a 12-h dark-light cycle (light from 07:00 to 19:00) at 22 ± 1°C with food and water provided ad libitum. Five postnatal 9- to 12-week-old male C57BL/6J mice (SLC, Shizuoka, Japan) were used.

### Preparation of Recording Electrodes

Each recording electrode for hippocampal LFP consisted of four tetrodes (diameter 17 μm, size 0007, and polyimide-coated platinum-iridium alloy (90/10%), California Fine Wire Company) with depths from the brain surface that were independently adjustable. The main body for fixing the substrate (EIB-36-PTB, Neuralynx) was three-dimensionally designed with 3D CAD Fusion360 (AUTODESK) and was formed with a photocurable 3D printer (Form2, Formlabs). The platinum coating was applied at the tips of tetrodes so that the impedances ranged from 150–300 kΩ.

### Surgery

Mice were anesthetized with 3% isoflurane inhalation. Under a concentration of 1.0– 1.5% isoflurane, electrode implantation was performed as follows. Anesthesia was confirmed by the lack of paw withdrawal, whisker movement, and eye-blink reflexes. The skin was subsequently removed from the head. A craniotomy (2.5 × 2.0 mm^2^) was performed, which was centered at 1.8 mm posterior to the bregma and 1.8 mm ventrolateral to the sagittal suture. Two screws (0.8 × 3.0 mm^2^; Muromachi Kikai Co., Ltd.) were embedded in the skull at the bilateral cerebellum until they reached the brain surface. The tips of the tetrodes were placed on the brain surface. One of the four electrodes was used as a reference by placing it in a shallow layer in which cortical firings were not recorded. The surface of the brain and the areas around the tetrodes were covered with Kwik-Sil Silicone Elastomer (World Precision Instruments). The skull surface was thinly covered with an adhesive (Super Bond C&B, Sun Medical) and was fixed on the skull using dental cement (Refine Bright, Yamahachi Dental Mfg., Co.). The screw used in this process served as a ground, and this ground and the ground of the board (EIB36-PTB, Neuralynx) were connected with a wire. A wire (A633, COONER WIRE) to record myoelectric potential was inserted into the trapezius muscles (muscles at the base of the head and neck). Finally, the ground wire and wire used to record the myoelectric potential were covered with dental cement. After the surgeries, electrodes were covered with a cap made with a hot-melt type 3D printer (UP Plus2, Sun Stella Co., Ltd.). Then, each mouse was returned to its cage.

### Adjusting the Recording Electrode Positions in the Hippocampus

After the operation, the electrode positions in the hippocampus were adjusted while the mice were in their home cages. Every screw for adjustment was tightened one turn every few minutes, which deepened the screw by 250 μm, until 1,000 μm from the brain surface was reached. With the waveforms monitored, each screw was tightened from 1/8 to 1/4 turn each time until the recording electrode reached a point just above the hippocampus. Then, each screw was loosened two turns and was left still for one day. Each screw was tightened by 1/4 turn each day until each electrode reached the target depth, and the maximum rotation angle of the screw was limited to 1/8 turn per day. The adjustment was stopped at the point at which large-amplitude ripples were observed.

### Electrophysiological Recording

The hippocampal LFP and trapezius myoelectric potential (MEP) were simultaneously recorded for five mice. After the recording electrodes in the hippocampus reached the desired depths, which was more than 14 days after surgery, the recording was performed with a data acquisition system (CerePlex Direct, Blackrock) for up to five days in the home cages of mice or a novel environment depending on the trial day, as shown in Table 1.

**Table 1.** Recording Environments. Mice #1, #2–4, and #5 were recorded for one day, four consecutive days, and five straight days, respectively. On the first and third days, experiments were performed in the home cages of mice. On the second, fourth, and fifth days, experiments were performed in a novel environment.

| Mouse # | Day 1 | Day 2 | Day 3 | Day 4 | Day 5 |
| --- | --- | --- | --- | --- | --- |
| #1 | Home | - | - | - | - |
| #2 | Home | Novel | Home | Novel | - |
| #3 | Home | Novel | Home | Novel | - |
| #4 | Home | Novel | Home | Novel |  |

**Table 1. Recording Environments.**
| #5 | Home | Novel | Home | Novel | Novel |
| --- | --- | --- | --- | --- | --- |

The sampling rate for recording the raw hippocampal LFP and trapezius MEP was 2 kHz. The raw data were recorded after applying a 500-Hz low-pass filter. Data were recorded under free-moving conditions and with food and water provided ad libitum. The novel environment was more spacious than the home cages, and it included three different objects. All recordings were performed under a 12-hour light-dark cycle with the dark period beginning at 19:00.

### Histochemical Verification of Recording Sites

After the recording experiments, the mice were anesthetized with urethane. Anesthesia was confirmed by the lack of paw withdrawal, whisker movement, and eye-blink reflexes. Overall, 25-μA currents were applied via electrodes in the hippocampus for 10 seconds to burn tissues at the recording sites. After the chest was dissected, ice-cooled phosphate-mM of KH_2_PO_4_) and 4% paraformaldehyde (PFA) phosphate buffer solution (4% PFA, in PBS) were perfused from the left ventricle. The head was cut and allowed to stand overnight. The brain was then removed and immersed in 4% PFA overnight and then immersed twice in a 30% sucrose/PBS solution overnight. The brain was snap-frozen on dry ice and stored at −80°C. Coronal sections with a thickness of 40 μm were prepared using a cryostat (CM3050 S, Leica). Each brain section was mounted on a microscope slide, stained with Cresyl Violet, and enclosed with a coverslip. Recording sites in the hippocampus were validated with records of the 3D coordinates of tetrodes, the number of electrodes, and the burn marks in coronal sections.

### Data Source

The identifiers used for the data are shown in Table 2.

**Table 2.**
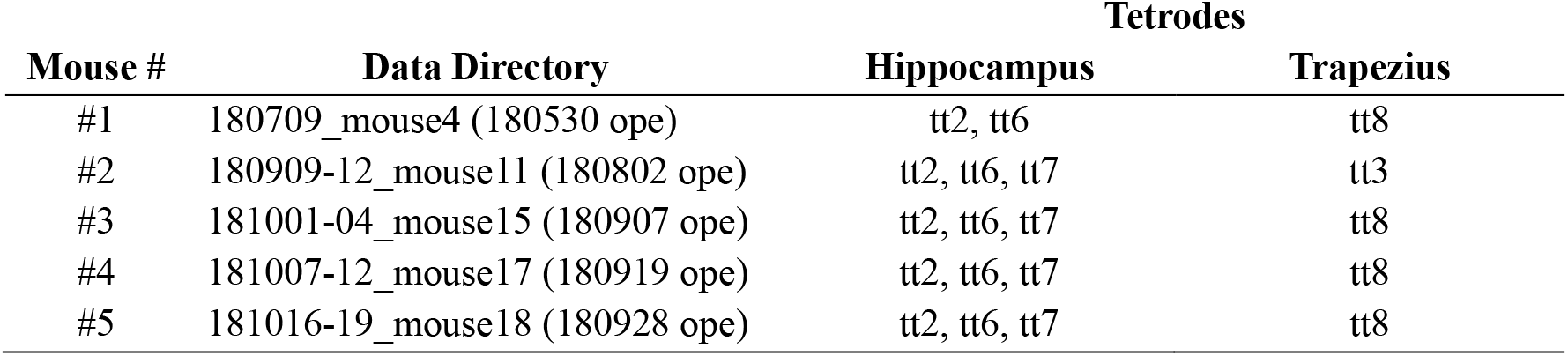
Data Source. The name of each data directory indicates the day of the surgery and the first day of the recording. To record hippocampal LFP, up to three tetrodes, or 12 electrodes, were inserted in the hippocampus.

### Tools for Data Analysis

MATLAB 2017b (MathWorks, Natick, MA, USA; https://www.mathworks.com/products/matlab.html) was used to read the recorded data from disks. For other analyses, Python 3.6.8 (https://www.python.org/) was used. Table 3 shows the explicitly imported Python packages.

**Table 3.** Explicitly Imported Libraries. The table shows the explicitly imported Python libraries used in this research, although there were other implicitly working packages and dependent packages.

| Library | Version | Project Page | Note |
| --- | --- | --- | --- |
| apex | - | https://github.com/NVIDIA/apex | - |
| cleanlab | 0.1.0 | https://github.com/cgnorthcutt/cleanlab | Northcutt et al., 2019 |
| cliffsDelta | - | https://github.com/neilernst/cliffsDelta | - |
| numpy | 1.17.0 | https://numpy.org | - |
| torch | 1.2.0 | https://pytorch.org | Also called as PyTorch |
| ranger | 9.3.19 | https://github.com/lessw2020/Ranger-Deep-Learning-Optimizer | - |
| ripple_detection | 0.1.7.dev0 | https://github.com/Eden-Kramer-Lab/ripple_detection | Kay et al., 2016 |
| statsmodels | 0.11.0 | https://github.com/statsmodels/statsmodels | - |
| scipy | 1.2.0 | https://www.scipy.org | - |
| scikit-learn | 0.21.2 | https://scikit-learn.org | - |

### Definition of Ripple Candidates

The recorded hippocampal LFP amplitude vector was downsampled from 2 kHz to 1 kHz using the imported function “scipy.signal.decimate,” in which an eighth-order Chebyshev type-one filter was used as an anti-aliasing filter. This process can be expressed by the pseudocode below.

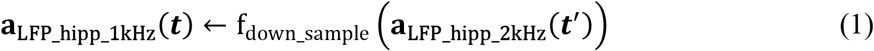

where ***a***_LFP_hipp_1kHz_(***t***) ∈ ℝ^T^ is the downsampled hippocampal LFP amplitude vector with a virtual sampling rate of 1 kHz, f_down_sample_(⋅) is the imported Python function “scipy.signal.decimate,” and ***a***_LFP_hipp_2kHz_(***t***′) ∈ ℝ^2T^ is the hippocampal LFP amplitude vector recorded at a sampling rate of 2 kHz. Then, the ripple band LFP was extracted by filtering the downsampled LFP series ***a***_LFP_hipp_1kHz_(***t***) with a bandpass filter for the ripple band (150–250 Hz) based on the pseudocode below.

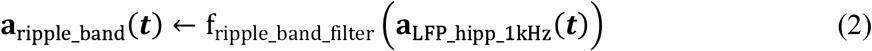

where ***a***_ripple_band_(***t***) is the extracted ripple band LFP amplitude vector and f_ripple_band_filter_ is the imported Python function “ripple_detection.core.filter_ripple_band.” By taking the square of the ripple band LFP amplitude [mV] at each time point, the ripple band LFP power [mV^2^] was calculated.

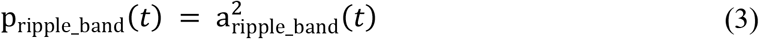

where p_ripple_band_(***t***) and a_ripple_band_(***t***) are the ripple band LFP power [mV^2^] and ripple band LFP amplitude [mV] at time *t*, respectively. The ripple band LFP power was smoothed over time using a 32-ms Gaussian kernel with the parameter σ = 4 ms. This was conducted with the imported Python function “ripple_detection.core.gaussian_smooth” as expressed in the following pseudocode:

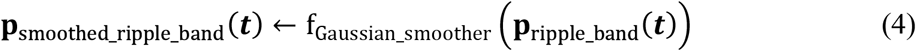

where ***p***_smoothed_ripple_band_(***t***) is a Gaussian-smoothed ripple band LFP power vector and f_Gaussian_smoother_(⋅) is the imported Python function “ripple_detection.core.gaussian_smooth” with the parameter σ = 4 ms. Then, the ripple band LFP magnitude [mV] at time *t* was defined as the root of the smoothed ripple band LFP power [mV^2^] at the corresponding time as follows:

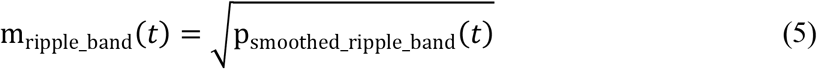

where m_ripple_band_(*t*) is the ripple band LFP magnitude [mV] at time *t*.

Ripple candidates were defined as LFP events with ripple band LFP magnitudes that continuously exceeded 1 SD for at least 15 ms. The defined onsets and offsets were redefined to the time points at which the ripple band LFP magnitude reached the mean value based on the imported Python function “ripple_detection.core.extend_threshold_to_mean,” as shown below.

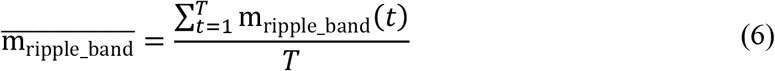

where *T* is the total sampling time for the electrode, which is calculated by multiplying the recording time [sec] by the virtual sampling rate of 1,000 [Hz]. Consequently, every ripple candidate was defined with a set of onset and offset times. As shown in Fig. 1, the normalized ripple band LFP magnitude at time *t* was calculated as follows:

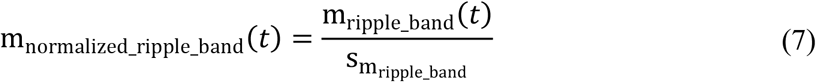

where m_normalized_ripple_band_(*t*) is the normalized ripple band LFP magnitude at time *t* and 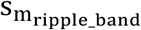 is the standard deviation of the ripple band LFP magnitude [mV] over time *t* as

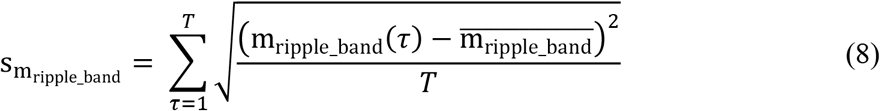

**Figure 1.**
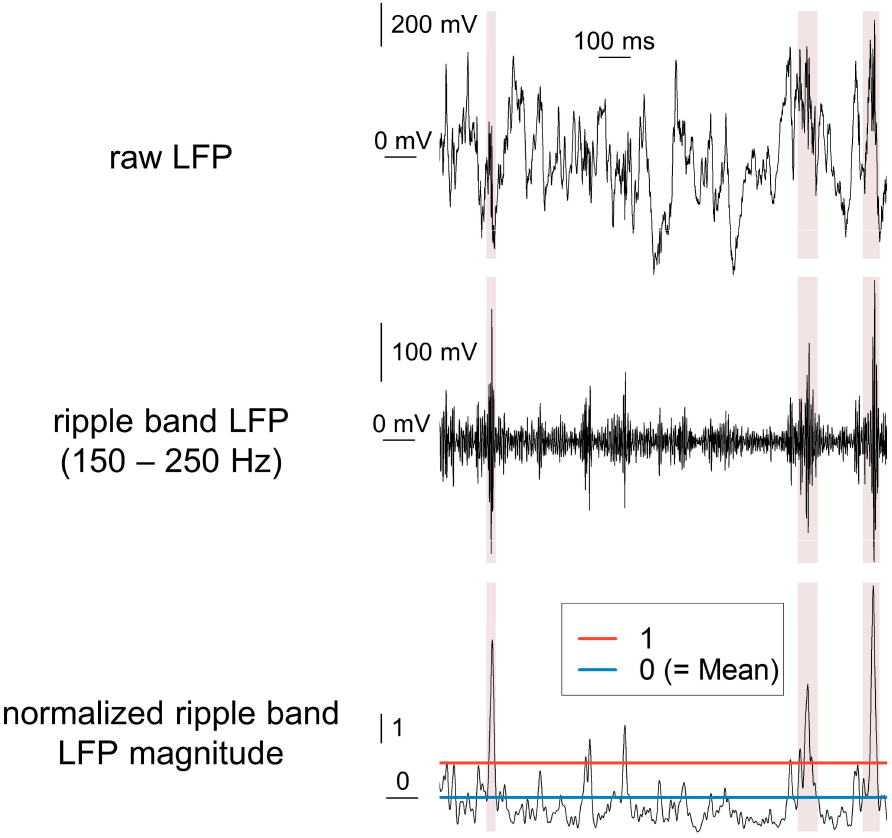
Selection of ripple candidates. *Top*: An LFP trace recorded from the CA1 pyramidal layer of the right hippocampus from a freely moving mouse. *Middle*: The corresponding 150–250 Hz-bandpass LFP trace as expressed in Equation (1&2). *Bottom*: The corresponding normalized ripple band LFP magnitude as defined in Equation (1–8). Superimposed red areas show the periods selected as ripple candidates.

### Definition of the “Three Variables” of Ripple Candidates

From every ripple candidate, the following “three variables” were calculated: the duration of each ripple candidate [ms], the normalized peak magnitude of each ripple candidate [a.u.], and the mean normalized magnitude of the MEP of the trapezius for each ripple candidate [a.u.] (Fig. 2, 4).

**Figure 2.**
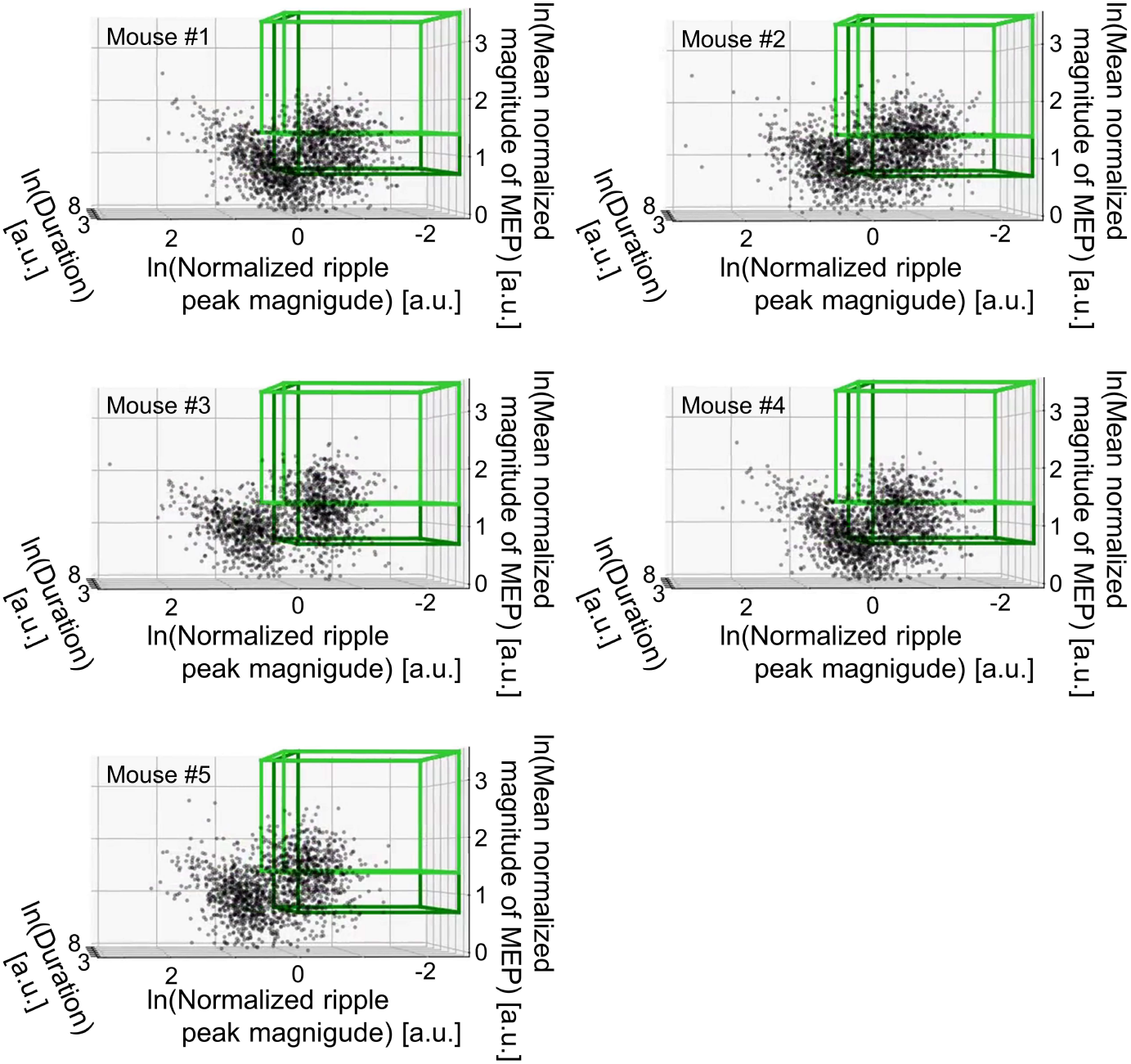
Three-dimensional visualization of ripple candidates. Ripple candidates were three-dimensionally plotted with a natural logarithm scale for the duration, mean normalized magnitude of the myoelectricity potential (MEP) of the trapezius, and normalized ripple peak magnitude (ripple band: 150–250 Hz) for each ripple (about the definitions, see Equations (9)–(22) in Materials and Methods). Each panel shows data for each mouse. Mice #1, #2–#4, and #5, data were continuously recorded for one, four, and five days, respectively. For visualization purposes, 0.20% and 0.05% of all ripple candidates recorded from mice #1 and #2–#5, respectively, were randomly sampled and plotted. The light and dark green cubes show the regions in which the ripple candidates are defined as “ripples” with the criteria of Fernández-Ruiz et al. (2019) and Kay et al. (2016); these cubes do not overlap.

The duration [ms] was defined as follows:

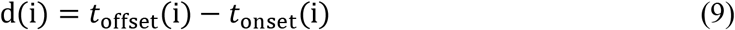

where d(i) is the duration [ms] of the i-th ripple candidate and *t*_offset_(*i*) and *t*_onset_(*i*) are the offset [ms] and onset [ms] of the i-th ripple candidate, respectively. Note that from the definition of ripple candidates (see the previous section), the ripple band LFP magnitude at both onsets and offsets was the mean of the ripple band LFP magnitude for each recording electrode.

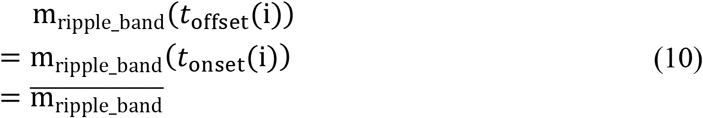

where m_ripple_band_(*t*) is the ripple band LFP magnitude [mV] at time *t*, as defined in Equation (5), and 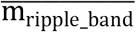 is the mean of the ripple band LFP magnitude for each recording electrode, as defined in Equation (6).

The normalized ripple peak magnitude [a.u.] was defined as follows:

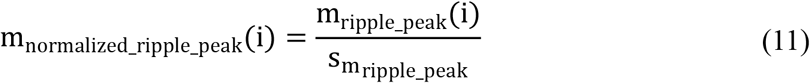

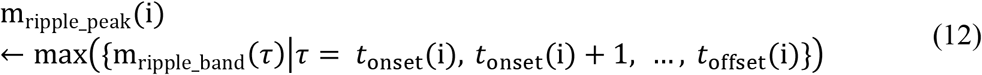

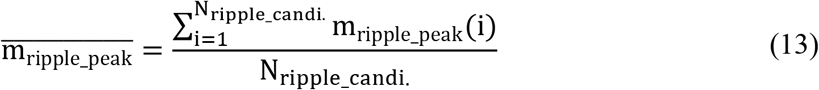

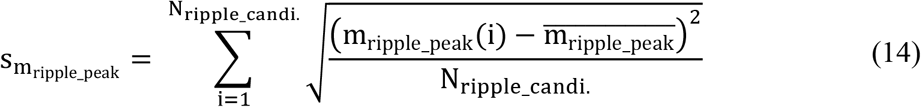

where m_normalized_ripple_peak_(i) is the normalized ripple peak magnitude [a.u.] for the i-th ripple candidate, m_ripple_band_(τ) is the ripple band LFP magnitude [mV] at time τ, as defined in Equation (5), and N_ripple_candi._ is the sample size of ripple candidates defined from the corresponding electrode.

The mean normalized magnitude of the MEP of the trapezius for ripple candidates [a.u.] was defined as follows. First, the magnitude of the MEP was obtained as the same as the pseudocode and Equations (1) and (3)–(5).

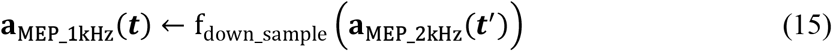

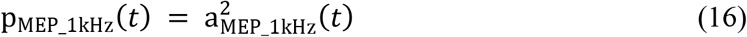

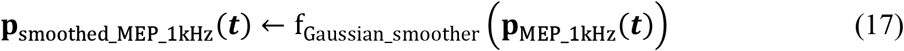

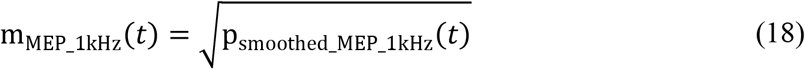

where ***a***_MEP_1kHz_(***t***) ∈ ℝ^T^ is the downsampled MEP amplitude vector with a virtual sampling rate of 1 kHz, f_down_sample_(⋅) is the imported Python function “scipy.signal.decimate,” ***a***_MEP_2kHz_(***t***′) ∈ ℝ^2T^ is the MEP amplitude vector recorded at 2 kHz, f_Gaussian_smoother_(⋅) is the imported Python function “ripple_detection.core.gaussian_smooth” with the parameter σ = 4 ms, and m_MEP_1kHz_(*t*) is the magnitude of the MEP at the time *t*. Finally, by averaging the magnitude of the MEP during a ripple and dividing the result by SD, we defined the normalized magnitude of the MEP for a ripple candidate [a.u.] as follows:

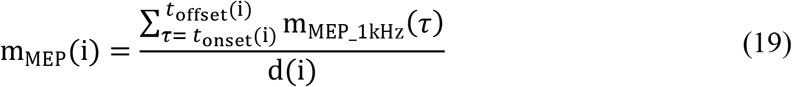

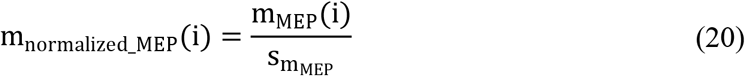

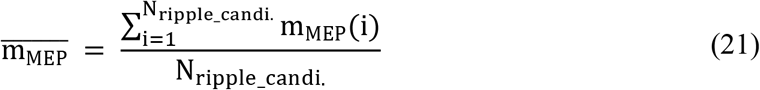

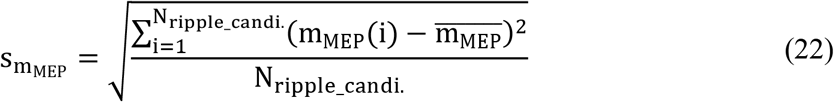

where m_MEP_(i) is the mean magnitude of the MEP of the i-th ripple candidate, m_normalized_MEP_(i) is the normalized mean magnitude of the MEP of the i-th ripple candidate, and N_ripple_candi._ is the sample size of the ripple candidates defined from the corresponding electrode.

### Relationship Between the Hippocampal LFP and Myoelectricity of the Trapezius

After being given a random initial sampling time ranging from 0–1,023 ms, hippocampal LFP data from each recording electrode were binned to consecutive 1,024-ms samples without overlaps. Additionally, the MEP of the trapezius during the corresponding periods was sampled.

In each 1,024-ms bin, the mean magnitude of the MEP was determined as shown below.

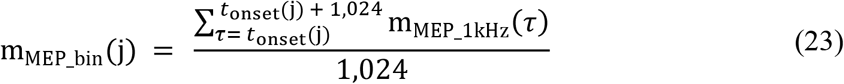

where m_MEP_bin_(j)  is the mean magnitude of the MEP [mV] of the j-th MEP sample, *t*_onset_(j) is the onset [ms] of the j-th time bin, and m_MEP_1kHz_(τ) is the magnitude of the MEP [mV] at time τ, as calculated from the pseudocode and Equations (15)–(18). Then, the mean magnitude of MEP was normalized as follows:

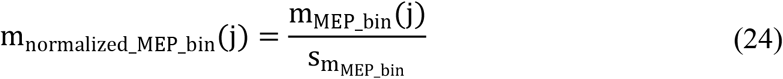

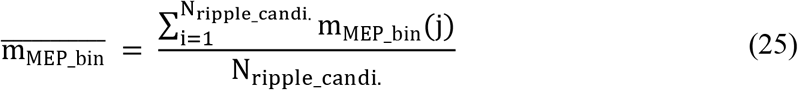

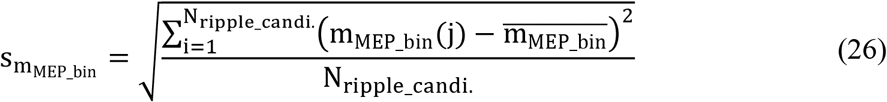

From the hippocampal LFP in each time bin, the FFT power [mV^2^ Hz^−1^] of f Hz component (f = 0, 1,…, 499) was obtained using the imported Python function “scipy.fftpack.fft.”

The Pearson correlation coefficients between the mean magnitude of the MEP [a.u.] and the FFT powers [mV^2^ Hz^−1^] of f Hz were calculated as follows:

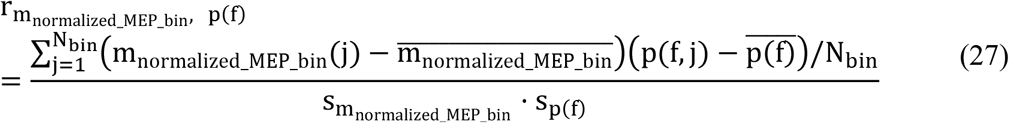

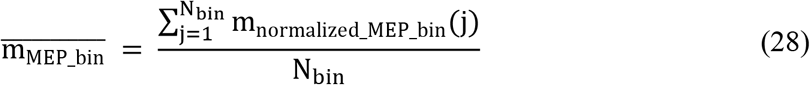

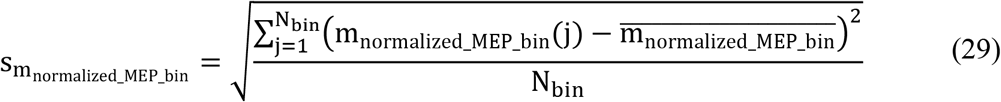

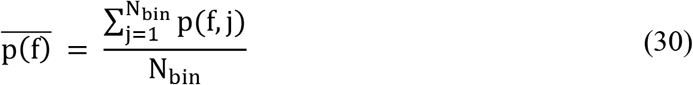

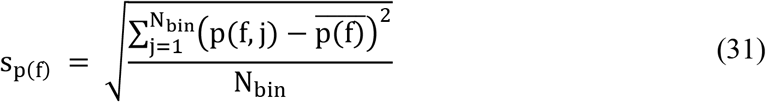

where 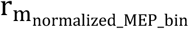, p(f) is the Pearson correlation coefficient [a.u.] between the normalized mean magnitude of the MEP [a.u.] and the FFT power [mV^2^] of the f Hz component (f = 0, 1,…, 499) of the hippocampal LFP for 1,024-ms bins, j is the index for time bins, N_bin_ is the total number of time bins, and p(f, j) is the FFT power [mV^2^ Hz^−1^] of the f Hz component of the j-th 1,024-ms hippocampal LFP sample (Fig. 3).

**Figure 3.**
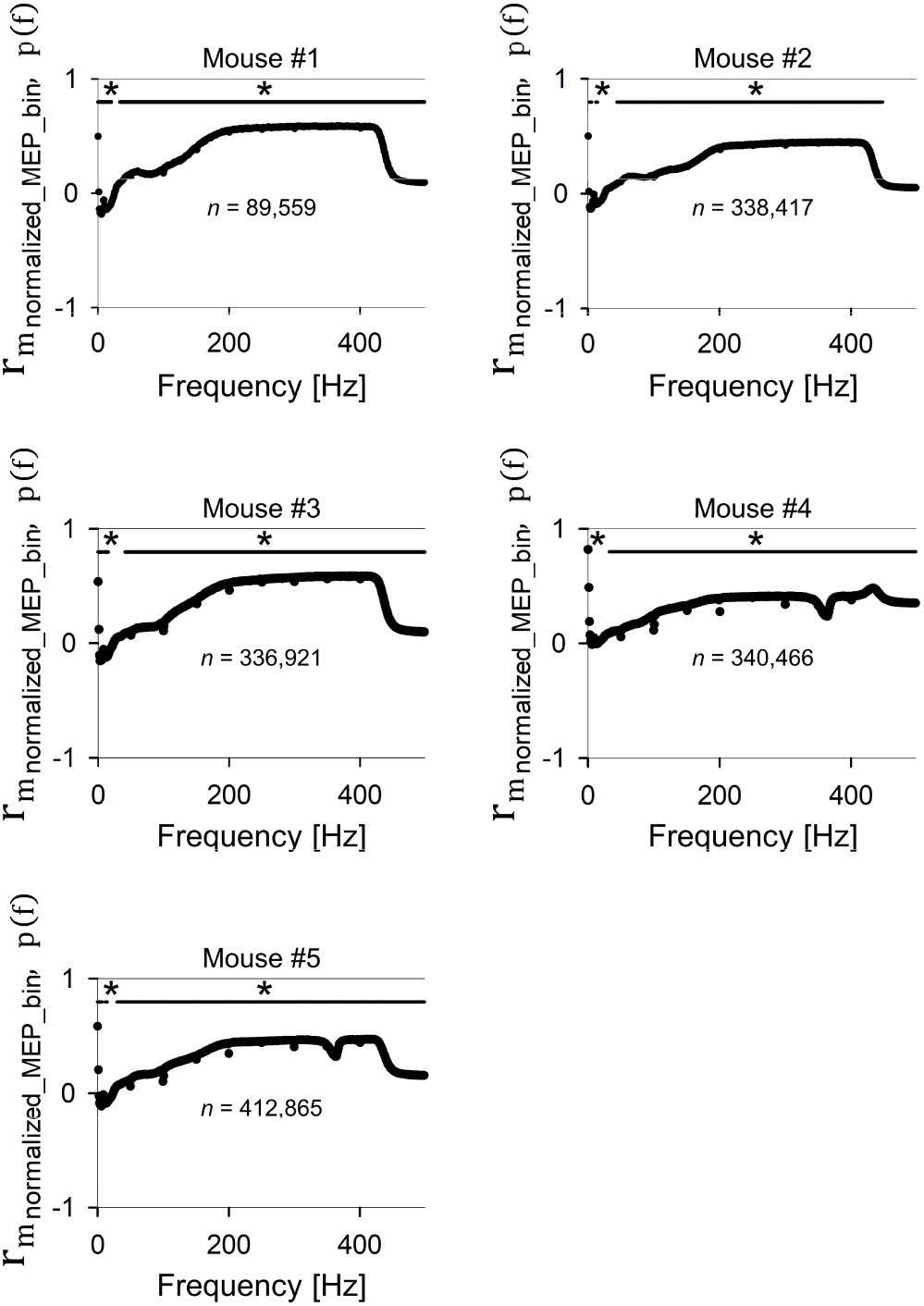
Relationship between the hippocampal LFP and animal movement. The Pearson correlation coefficients between the normalized mean of MEP of the trapezius (m_normalized_MEP_bin_(j) [a.u.]; j is the index for 1,024-ms bin) and the FFT power of the f-Hz component of the hippocampal LFP (p(f, j) [mV^2^ Hz^−1^]; j is the index for 1,024-ms bin) plotted against frequencies [Hz] (see Equations (23)–(31) in Materials and Methods). Each panel shows data for each mouse. The correlations from 44–448 Hz were statistically significant in all mice (*p < .05; the test for no correlation, ns, for the 1,024-ms time samples in each panel).

### Partial Correlation

The partial correlation between variables P and Q, controlling for the influence of variable *R*, can be expressed as r_PQ_R_, as shown as below.

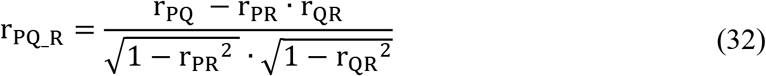

where r_PQ_, r_PR_, and r_QR_ are the Pearson correlation coefficients between the index variables.

### Construction of a Deep Convolutional Neural Network (CNN)

Based on Fawaz et al. (2019), as one of the best models for time series classification, we chose ResNet (He et al., 2015), a deep CNN. We implemented ResNet with six minor revisions using PyTorch 1.2.0 (https://pytorch.org/), a framework for machine learning. We call the modified model “our CNN” in this paper (Fig. 6). Specifically, the six modifications were as follows. (i) The first filter length in each “block” was shortened from 8 to 7. (ii) The activation function was changed from the ReLU function (Hahnloser et al., 2000) to the LeakyReLU function (Maas et al., 2013), and the differential coefficient in the negative region was 0.1. (iii) The number of filters in every convolutional layer was quadrupled (e.g., 64 to 256). (iv) Two fully connected layers and a softmax layer were added at the end in the same order as listed. These two layers functioned as a stochastic classifier for two-class classification tasks. Dropout (Srivastava et al., 2014) was applied in the first fully connected layer to prevent overfitting. Note that in our model, dropout was only applied on the downstream side of the batch normalization (Ioffe and Szegedy, 2015) layers because it was reported that the coapplication of dropout and batch normalization functions might lead to incongruity in results (Li et al., 2019). (v) The model performed mixed-precision training based on the imported Python module “apex”; the model was able to process data not only in a 32-bit floating format but also in a 16-bit floating format. (vi) The model was replicated on four GPUs (ASUS GeForce GTX 1080 TI 11GB Turbo × 4) with the imported Python module “torch.nn.parallel,” and a multi-GPU parallel computing model was formed.

### Isolating and Trimming Each Ripple Candidate as the CNN Inputs

Our CNN requires an input vector to be a fixed length. We fixed the input length at 400 ms for a sampling rate of 1 kHz. Thus, the i-th input vector was expressed as **x**_*i*_ ∈ ℝ^400^. If an input vector **x**_*i*_ ∈ ℝ^400^ includes multiple ripple candidates, the output may be unclear. Therefore, the following two steps were performed for each ripple candidate: (i) isolating the candidate and (ii) setting the time length to 400 ms by either zero-padding or trimming.

First, each ripple candidate **c**_*i*_ ∈ ℝ^d(i)^ was isolated from the onset to the offset as follows.

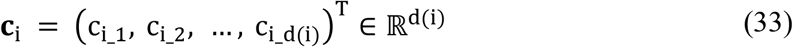

where d(i) is the duration [points] (or [ms]) of the i-th ripple candidate as defined in Equation (9). When the duration of a candidate was less than 400 ms, the candidate was padded with zeros to be extended symmetrically from the middle time (i.e., zero-padding). If the duration was 400 ms or longer, the LFP signal around the middle time was extracted (i.e., trimming). The zero-padding and trimming functions are as follows:

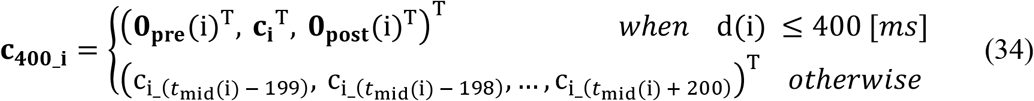

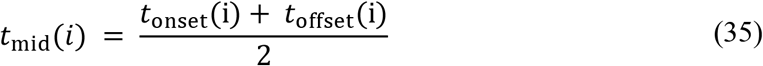

where ***c***_400_**i**_ ∈ ℝ^400^ is the 400-ms padded or trimmed data sequence derived from the i-th ripple candidate ***c***_***i***_  ∈ ℝ^*d(i)*^; d(i) is the duration [ms] for the i-th ripple candidate; 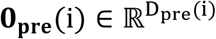 and 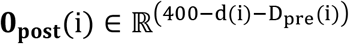 are zero vectors to pad; D_pre_(i) is defined as 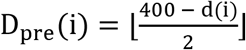; *t*_offset_(i) and *t*_onset_(i) are the offset and onset [ms] of the i-th ripple candidate, respectively; and c_i_*t*_ is the hippocampal LFP amplitude [mV] at time *t*, which is also a component of the i-th ripple candidate ***c***_i_.

### Data Splitting

Data splitting was performed according to the ID numbers allocated to mice to estimate matrices on unseen data. The following ten subdatasets were defined. Subdatasets SD1*, SD2*, SD3*, SD4*, and SD5* were defined as datasets that “only included” the data from mice #1, #2, #3, #4, and #5, respectively. Subdatasets SD1-, D2-, D3-, D4-, and D5- were defined as datasets that “excluded” the data from mice #1, #2, #3, #4, and #5, respectively.

### Confident Learning to Define Ripples

To define ripples from ripple candidates, we conducted Confident Learning with the imported Python module “cleanlab” (Northcutt et al., 2019; https://github.com/cgnorthcutt/cleanlab). Confident Learning was applied to process each dataset via a process similar to the 5-fold cross-validation method. First, in each dataset, ripple candidates were randomly divided into five folds (e.g., “fold A–E”) without overlaps. Then, the labels of one of the five folds (e.g., “fold A”) were estimated using the remaining folds (e.g., “folds B–E”) as training data.

Thus, label estimation by Confident Learning was performed 50 times in total (ten datasets × five patterns according to the folds). Each ripple candidate was tagged with five estimated labels depending on the datasets considered to run Confident Learning. For example, the ripple candidate acquired from mouse #1 was included in the subdatasets SD1*, SD2-, SD3-, SD4-, and SD5-. Then, the candidate was independently assigned five labels for the five datasets.

### Optimization of our CNN to Define Ripples in Confident Learning

Our CNN requires the input length to be fixed. We set the input length to 400 [points] (or [ms] in our case since the virtual sampling rate was 1 kHz). The mini-batch size was set to 6,000. The cross-entropy loss function was used. The learning rate was fixed to 1.0 × 10^−3^. Training was performed in a supervised manner for two epochs. At the beginning of each epoch, the order by which samples {***c***_400_**i**_} (i = 1, 2,…) were fed into the CNN was shuffled. As the optimizer, the imported Python class “ranger. Ranger” was used based on combining two mechanisms: RAdam (Tong et al., 2019) and Lookahead (Zhang et al., 2019). The dropout rate of the first fully connected layer from the input side was set to 0.5 during the training stage.

Additionally, during the inference stage, the dropout value was set to zero, and batch normalization was performed in inference mode. The forward path of our CNN used to define ripples can be expressed by the following pseudocode and equation set:

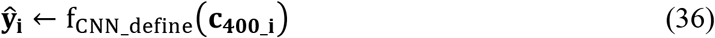

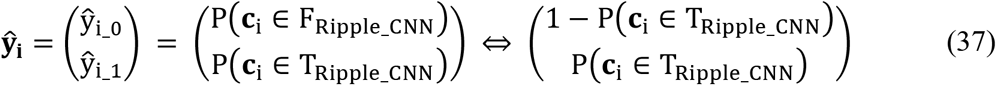

where ***c***_400_**i**_ ∈ ℝ^400^ is the 400-ms data sequence derived by padding or trimming the i-th ripple candidate ***c***_**i**_  ∈ ℝ^*d(i)*^, as defined in Equation (31); f_CNN_define_(⋅) is the forward path used by our CNN to define ripples; F_Ripple_CNN_ and T_Ripple_CNN_ are the labels indicating false and true ripples estimated by our CNN, respectively; and P(***c***_**i**_ ∈ T_Ripple_CNN_) is the probability that candidate c_i_ is defined as a true ripple by our CNN.

Based on the output 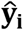, estimated labels (= T_Ripple_CNN_ or F_Ripple_CNN_) were allocated for each ripple candidate ***c***_**i**_ by setting a decision threshold (= d_threshold_define_) as follows:

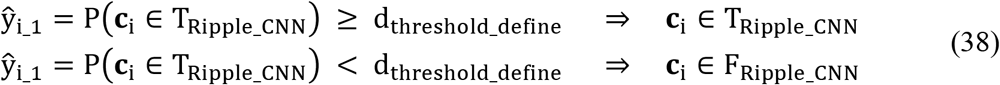

### Optimization of our CNN for Ripple Detection

Our CNN requires the input length to be fixed. We set the input length to 400 [points] (or [ms] in our case since the virtual sampling rate was 1 kHz). The mini-batch size was set to 1,792. The modified cross-entropy loss function was used, as expressed in the following section titled “Balancing the Loss Function for an Imbalanced Dataset.” The learning rate was exponentially reduced with the training iteration number from the initial value of 1.0 × 10^−3^ to the final value of 1.0 × 10^−4^ based on the imported Python class “torch.optim.lr_scheduler.StepLR” (the exponential learning decay). The training was performed for three epochs in a supervised manner on labels acquired by an unsupervised Gaussian mixed model. At the beginning of each epoch, input signals {***s***_**i**_} (i = 1, 2,…) were resampled by the random systematic sampling method over time, and the order of inputs for the CNN was shuffled. Note that for simplicity, 400-ms raw LFP signals that included more than any part of two ripples, tagged as T_Ripple_CNN_, were excluded from our analyses. As the optimizer, the imported Python class “ranger.Ranger” was used. The dropout rate of the first fully connected layer from the input side was set to 0.5 only during the training stage. During the inference stage, the dropout value was set to zero, and batch normalization was performed in inference mode. The forward path of our CNN used to detect ripples can be expressed by the following pseudocode and equation.

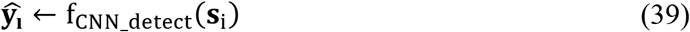

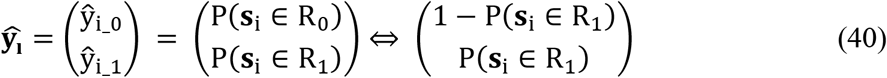

where ***s***_**i**_ ∈ ℝ^400^ is the i-th 400-ms raw LFP signal, f_CNN_detect_(⋅) is the forward path of our CNN used to detect ripples, 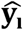 is the i-th output vector, and R_0_ and R_1_ are the labels and indicate that ***s***_**i**_ includes zero ripples or just one ripple, respectively.

Based on the output 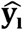, predicted labels 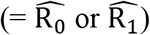 were allocated for each 400-ms raw LFP signal ***s***_**i**_ (i = 1, 2,…) by setting a decision threshold (= d_threshold_detect_) as follows:

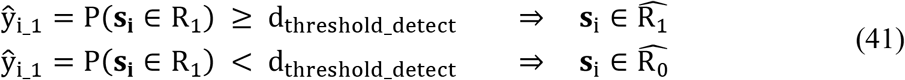

### Balancing the Loss Function for an Imbalanced Dataset

In the ripple detection task, the sample sizes of the “ripple-not-including” group (labeled R_0_) and “one-ripple-including group” (labeled R_1_) were imbalanced; the ripple-not-including group (R_0_) was more than 100 times larger than the one-ripple-including group (R_1_). To reflect the loss from the minority group (R_1_) during training, we modified the cross-entropy loss function in a cost-sensitive learning framework (Kukar et al., 1998). The cost function for the k-th training loop using the original cross-entropy loss function, J(θ)_k_, is written as follows:

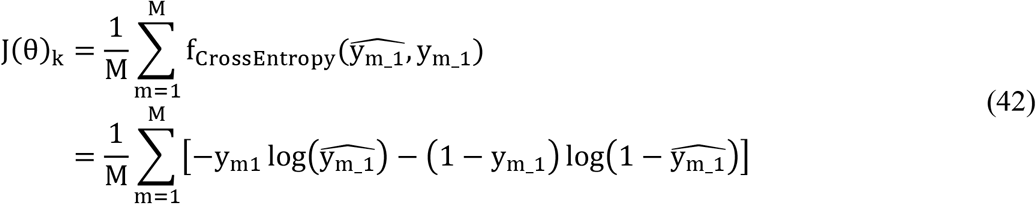

where M is the mini-batch size, m is the index of the samples in a mini-batch, 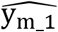 is the predicted probability for the m-th sample in the mini-batch to belong to the ripple-not-including group (R_1_) based on Equation (38), and y_m_1_ is the ground truth label (0: ripple-not-including group; 1: one-ripple-including group) for the m-th sample in the mini-batch. To address the sample size imbalance among classes, we modified the original cross-entropy loss function J(θ)_k_ in Equation (39) to J′(θ)_k_ by weighting the losses from each class considering the encounter rate for our CNN as follows:

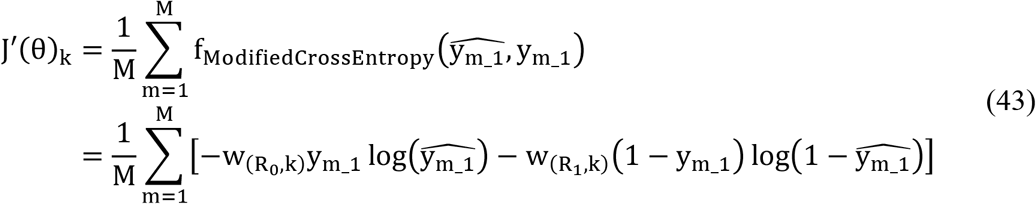

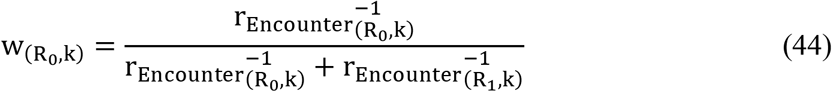

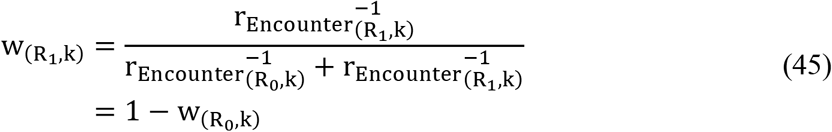

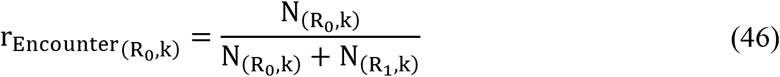

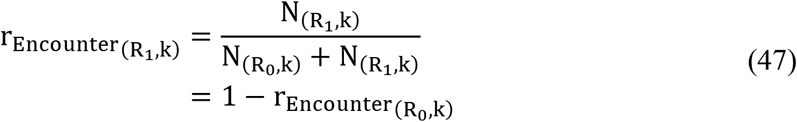

where J′(θ)_k_ is the modified cost function of the k-th training loop using the weighted cross-entropy loss function; 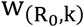 and 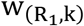 are the weights multiplied by the losses from the ripple-not-including group and the one-ripple-including group during the k-th training loop, respectively; 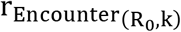 and 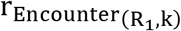 are the encounter rates of the ripple-not-including group and the one-ripple-including group, respectively; and 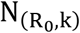 and 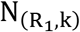 are the cumulative sample sizes of the ripple-not-including group and the one-ripple-including group through the k-th training loop, respectively.

## Results

### Thresholds in Existing Ripple Detection Methods and the Defined “Ripples”

Existing ripple detection methods first define ripples by selecting ripple candidates. Second, they sort the candidates by setting thresholds for features such as the ripple band RMS magnitude or power of each candidate, the duration of each candidate, and the animal head speed for each candidate. However, such thresholds can be arbitrarily set by analysts. There were no consistent values or acceptable ranges for the thresholds among the research groups. Here, we examined the effects of differences in the thresholds on defined “ripples.” Specifically, we used hippocampal LFP data and MEP data from the trapezius; these data were simultaneously recorded over up to five consecutive days from five mice.

First, ripple candidates were defined based on the definition of Kay et al. (2016) (Fig. 1; see Materials and Methods) with the following two modifications. First, the threshold of the peak magnitude of a ripple band was reduced from 2 SD to 1 SD, and second, the process of sorting ripple candidates using animal head speed was removed. These two modifications were implemented to include more true latent ripples in ripple candidate sets.

To define ripples from the candidates, we used the MEP of the trapezius and not the animal head speed for the following reasons. Existing methods define ripples only when the animal head speed is less than a threshold (e.g., 4 cm/s). Generally, vibrations or impacts on recording electrodes generate noise in recorded signals. Thus, setting a threshold for animal head speed avoids noise at times when detecting ripples is difficult due to physical activities. However, the reliability of animal head speed as a variable is relatively low compared to that of the ripple duration (e.g., < 50 ms), mainly because of the video-capturing limitations in timely and the spatial resolution (e.g., ≤ 60 fps and ~50 MPs); additionally, the MEP has higher time and amplitude resolutions (e.g., at a sampling rate of 20 kHz and for 16-bit signed integer data). Thus, we chose not animal head speed but MEP as an indicator of the noisy LFP period associated with animal movements.

First, from each ripple candidate, the following “three variables” were calculated: the duration [ms], normalized peak ripple magnitude [a.u.], and mean normalized magnitude of the MEP of the trapezius for a ripple candidate [a.u.] (see Materials and Methods). Additionally, “three ln-variables” were calculated by taking the natural logarithm of each of the “three variables.” Next, each ripple candidate was plotted in a 3D space with the “three ln-variables” as the axes (Fig. 2). In Fig. 2, detecting ripples with one of the existing methods (Ramirez-Villegas et al., 2015; Karlsson and Frank, 2009; Kay et al., 2016; Fernández-Ruiz et al., 2019; Shin et al., 2019; Hannah et al., 2019) is equal to establishing a specific cube in the 3D space in parallel with the axes and selecting the ripple candidates in the cube as ripples. For example, the dark and light green cubes in Fig. 2 are constructed based on the thresholds used by Kay et al. (2016) and Fernández-Ruiz et al. (2019), respectively. Here, we assumed that the mean normalized magnitude of the MEP of the trapezius for a ripple candidate [a.u.] is directly proportional to the animal head speed in the same time resolution with the corresponding research and the corresponding conversion equation is as follows.

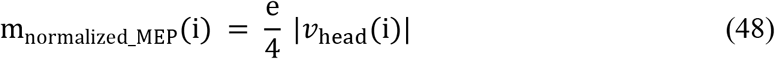

where m_normalized_MEP_(i) is the normalized mean magnitude of the MEP [mV] of the i-th ripple candidate, as defined in Equation (20), e is Napier’s constant, and *v*_head_ is the mean animal head speed [cm/s] during the i-th ripple candidate. As a result, the ripple candidates included in the cubes corresponding to the methods of Kay et al. (2016) and Fernández-Ruiz et al. (2019) were not consistent. As graphically shown in Fig. 2, it was revealed that “ripples” are sensitive to the thresholds that must be set in the existing detection methods.

### Relationship Between the Hippocampal LFP and Animal Movements

As mentioned in the previous section, screening ripple candidates using the animal head speed in the existing ripple detection methods leads to poor ripple detection due to noise during animal movement. Thus, we use the mean normalized magnitude of the MEP for screening. If these two assumptions are valid, inversely, the larger the mean normalized magnitude of the MEP is, the larger the amplitude of the ripple band (150– 250 Hz) LFP should be. We checked this hypothesis with our data.

The Pearson correlation coefficients between the mean magnitude of the MEP [a.u.] and the FFT power at f Hz [mV^2^ Hz^−1^] (f = 0, 1,…, 499) were calculated for 1,024-ms samples (Fig. 3; see Materials and Methods, especially Equations (23)–(31)). In all five mice, at 44–448 Hz [mV^2^ Hz^−1^], the correlations were statistically significant (ps < .05, the test for no correlation). This result can be interpreted as follows: while mice moved around, there were transient neuronal activities in the hippocampus. However, this result provided supporting evidence for the hypotheses discussed earlier: the recorded hippocampal LFPs are contaminated by noise caused by animal movements, and the mean normalized magnitude of the MEP [a.u.] is a useful barometer for noisy periods.

### Defining Ripples Based on Gaussian Mixed Clustering

As considered in the previous section, the existing detection methods are designed to exclude false ripples from ripple candidate sets. The sorting process can be optimized. In existing methods, although each ripple candidate is associated with the “three variables” determined at the same time, the threshold of each variable is individually determined; that is, the multivariate nature of these variables is not considered. In general, when trying to separate multivariate samples into two classes, an appropriate multivariate model yields better outcomes than individually applying univariate models. Here, the higher the correlation among variables is, the more dominant the positive effect of a multivariate model (Ichihara, 1990). Therefore, we assumed that the multivariate nature of the “three ln-variables” should be considered in defining hippocampal ripples.

Thus, we first determined the correlations among the “three ln-variables.” The partial correlation coefficients among these were calculated to exclude a possible spurious correlation (Table 4). For all three patterns, the partial correlations were statistically significant (ps < .05, multiple comparison using the test for no correlation with Bonferroni correction). These results suggest that the process used to distinguish false ripples from ripple candidates can be optimized using a multivariate approach.

**Table 4.** Partial correlations among the “three ln-variables” of ripple candidates. The partial correlation between variables P and Q, controlling for the influence of variable R, or r_PQ_R,_ is defined in Equation (32). The table shows the partial correlations among the “three ln-variables” of ripple candidates: ln(duration) [a.u.], ln(normalized ripple peak magnitude) [a.u.], and ln(mean normalized magnitude of MEP) [a.u.] for each ripple candidate. Each pair of the three “ln-variables” was correlated (ps < .05, multiple comparison using the test for no correlation with Bonferroni correction).

|  |  |  |  |
| --- | --- | --- | --- |
| ln(duration) | X | Z | Y |
| ln(normalized peak magnitude) | Y | X | Z |
| ln(mean normalized magnitude of MEP) | Z | Y | X |
| $r_{XY_Z}$ | $0.46 \pm 0.02$ | $0.20 \pm 0.03$ | $-0.37 \pm 0.14$ |

Next, we defined ripples by using the multivariate nature of the “three ln-variables.” First, the histograms of the ln-variables were plotted (Fig. 4). From the histograms, we hypothesized that each distribution of the variables could be approximated as the sum of two Gaussian functions. Thus, with the assumption that ripple candidates should be divided into two clusters in the 3D space with axes set based on the three ln-variables, we separated the ripple candidates into two clusters by using a Gaussian mixed model (GMM; Fig. 5). As mentioned earlier, true ripples are expected to be included in one cluster, the centroid of which is smaller in the dimension of the mean normalized magnitude of the MEP. We named this cluster “Cluster T.” The other cluster, “Cluster F,” is expected to include false ripples. We labeled the ripple candidates assigned to Cluster T and Cluster F as T_Ripple_GMM_ and F_Ripple_GMM_, respectively.

**Figure 4.**
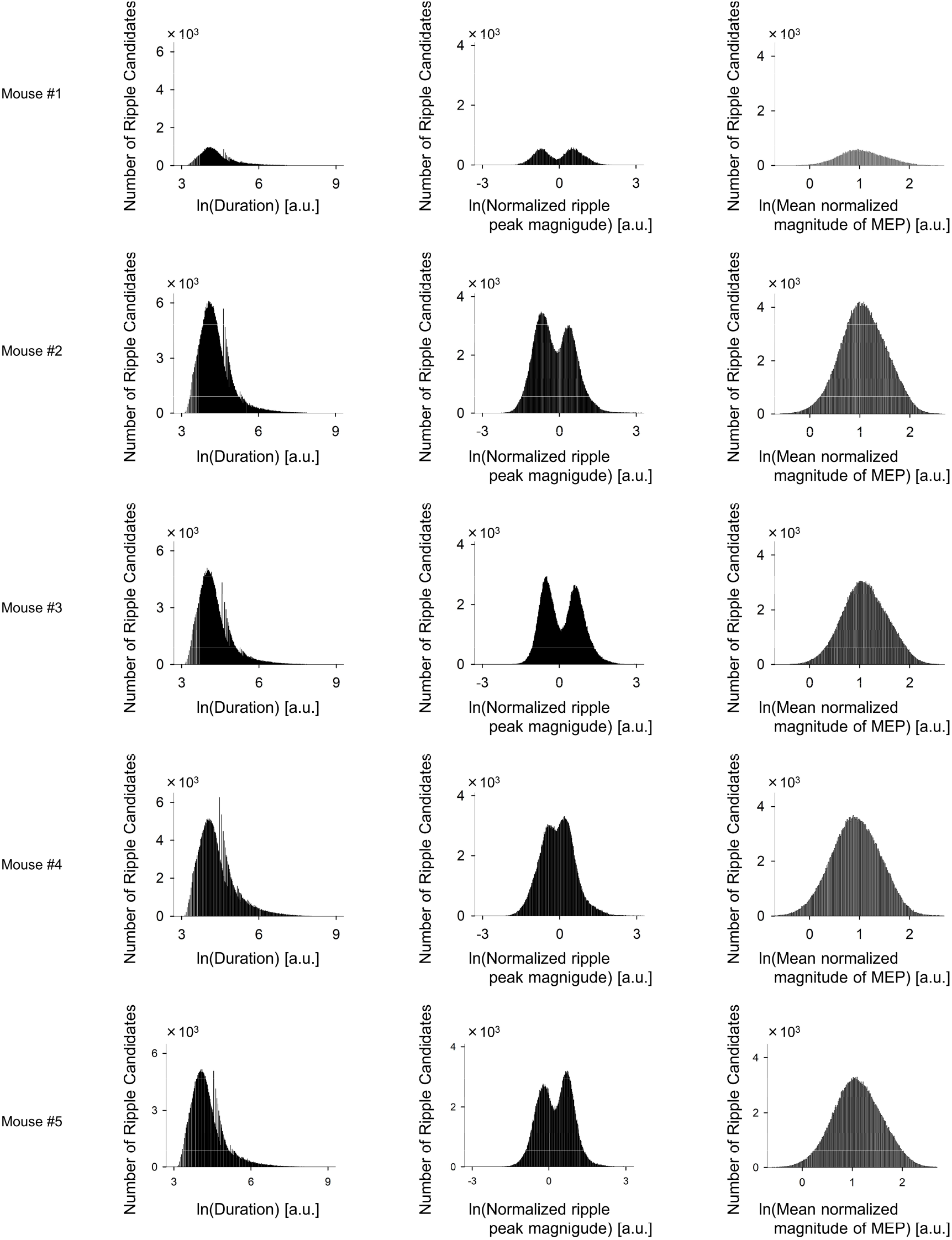
Distributions of the three parameters of the ripple candidates. Histograms of the ln(duration), ln(normalized ripple peak magnitude), and ln(mean normalized magnitude of the myoelectric potential (MEP) of the trapezius) for each ripple candidate. Each panel shows data for each mouse. We postulated that each distribution could be approximated as the sum of two Gaussian functions.

**Figure 5.**
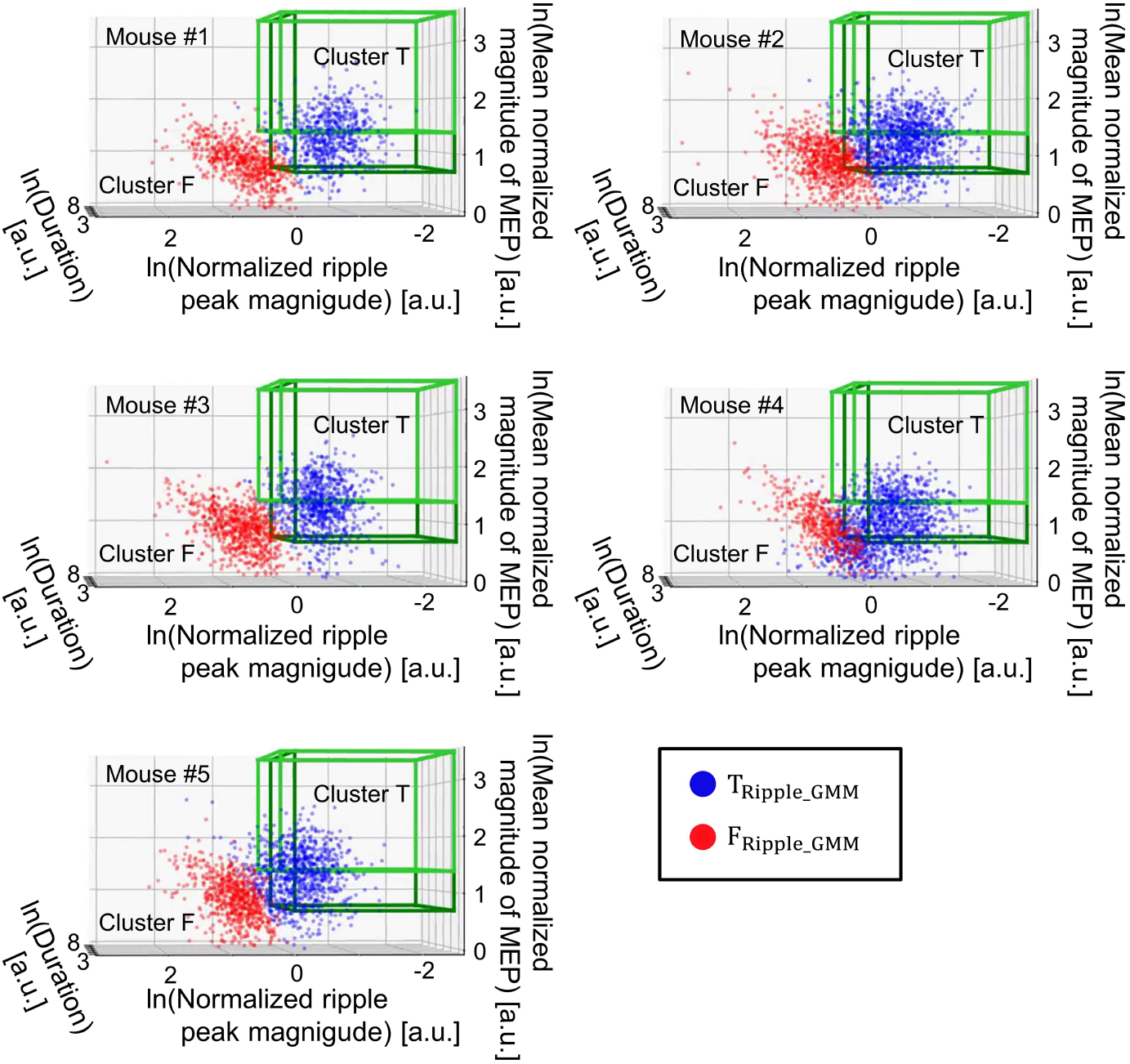
Ripples labeled using Gaussian mixed model clustering. Each plot in the 3D space shows true ripple-like events (*Blue*; tagged as T_Ripple_GMM_) and false ripple-like events (*Red*; tagged as F_Ripple_GMM_) defined with Gaussian mixed model clustering. Note that the axes of the 3D space coincide with the variables used in Gaussian clustering: ln(duration) [a.u.], ln (normalized ripple peak magnitude) [a.u.], and ln(normalized mean magnitude of the MEP of the trapezius) [a.u.]. The data cloud as a whole (T_Ripple_GMM_ and F_Ripple_GMM_) and light and dark green cubes are the same as those in Fig. 2. Note that the decision boundary determined by Gaussian mixed model clustering is a hyperplane in this 3D space.

### Defining Ripples by Using Our CNN in a Weakly Supervised Manner

Each ripple candidate was labeled with T_Ripple_GMM_ and F_Ripple_GMM_. However, the labels contained errors, or noise was present. In particular, GMM clustering uses only limited information from ripple candidates. The inputs for GMM clustering were the three ln-variables: ln(the duration of each ripple candidate) [a.u.], ln(the normalized peak magnitude of each ripple candidate) [a.u.], and ln(the mean normalized magnitude of the MEP of the trapezius for each ripple candidate) [a.u.]. The former two variables are predefined features extracted from each ripple candidate, and the final variable provides information about the noises associated with animal activities, as discussed earlier. The dimensionality reduction into the three ln-variables could not be appropriate.

Moreover, label errors can be found and mitigated by Confident Learning. Specifically, the imported Python module “cleanlab” (Northcutt et al., 2019) acts as an arbitrary classifier to estimate the latent true labels without hyperparameters. We conjectured that “noisy” labels, T_Ripple_GMM_ and F_Ripple_GMM_, could be “cleaned” by using the module with a CNN and a raw LFP input, because CNNs can search for better local features in original signals. The cleaned labels T_Ripple_GMM_ and F_Ripple_GMM_ are obtained from using optimal information about waveforms to distinguish true and false ripples, which is difficult for humans. Therefore, we calculated the cleaned labels T_Ripple_GMM_ and F_Ripple_GMM_ as follows.

First, we constructed a deep CNN (Fig. 6; see Materials and Methods). We simply call this model “our CNN” in this paper. Because only a fixed-length input (e.g., 400 ms at a sampling rate of 1 kHz in our case) can be used with this CNN, we isolated each ripple candidate and converted it to a 400-ms data series (Fig. 7; see Materials and Methods).

**Figure 6.**
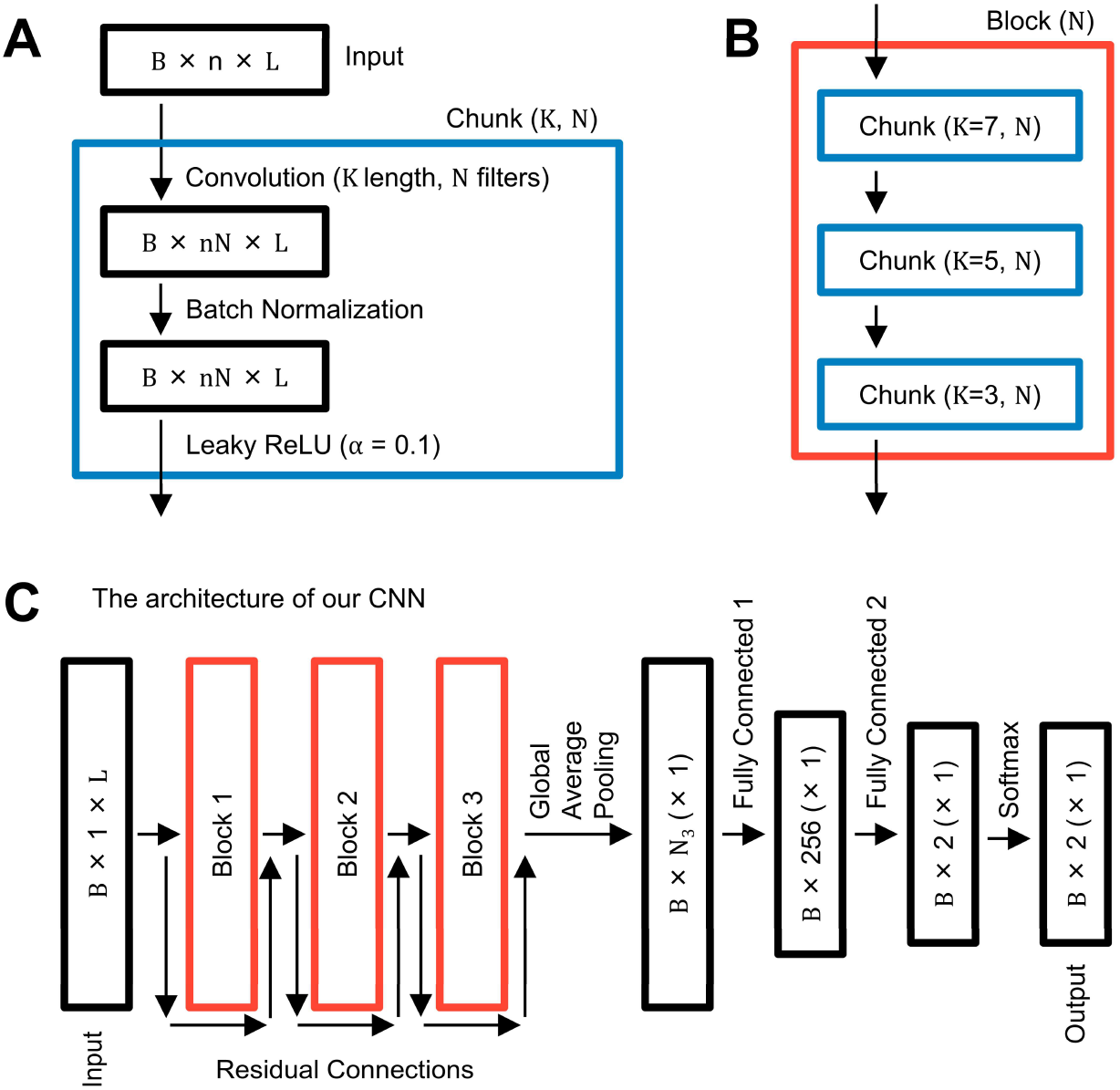
The structure of our CNN. **A.** A chunk (*Blue*) is the smallest unit in our CNN. A chunk has two parameters: K and N. N × convolution filters, each of which was of length K ([ms] in our usde case), were installed in the first layer. B shows the mini-batch size and the sample size used in training loops to accelerate the computing process. In batch normalization (Ioffe and Szegedy, 2015), the filtered traces are normalized and perturbed. Then, the output is activated by the Leaky ReLU activation function (Maas et al., 2013) with a differential coefficient of 0.1 in the negative region (α = 0.1). **B.** A block (*Red*) is the second smallest unit, and it includes three chunks with the parameters K = 7, 5, and 3. **C.** The whole structure of our CNN. To extract local features from input signals, blocks 1–3 were connected not only through the convolution pathway but also through residual connections (this model is called ResNet in general). Global average pooling was performed to “squash” the time dimension of the extracted features. Fully connected layers were adopted to add weights to the convolutional filters of the last layer and optimally use the extracted features for classification tasks.

**Figure 7.**
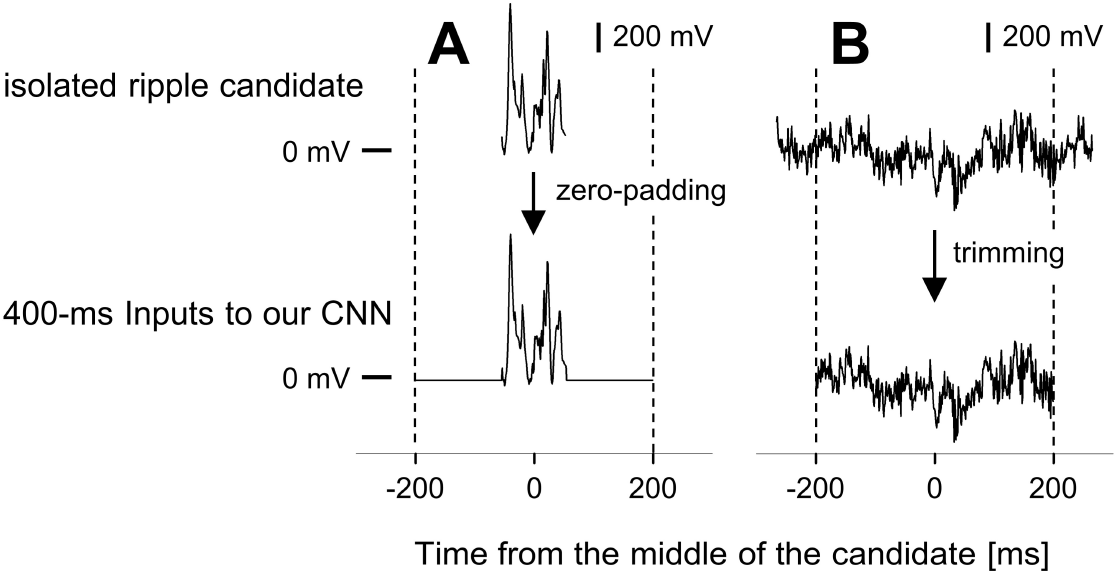
Isolating and trimming each ripple candidate as an input into our CNN. **A.** When a ripple candidate was 400 ms or less, it was extended with symmetrical zero-padding to 400 ms. **B.** Otherwise, information in the range of [-200 ms, +200 ms) from the middle time of a candidate was extracted (see Equation (34) in Materials and Methods). These synthesized 400-ms data series were used as inputs to our CNN to define ripples (see Equations (36)–(38) in Materials and Methods).

Next, data splitting was performed in Confident Learning (Fig. 8). The splitting aims to determine whether the different datasets influence label definition with Confident Learning and prevent “data leakage” in the subsequent ripple detection experiments.

**Figure 8.**
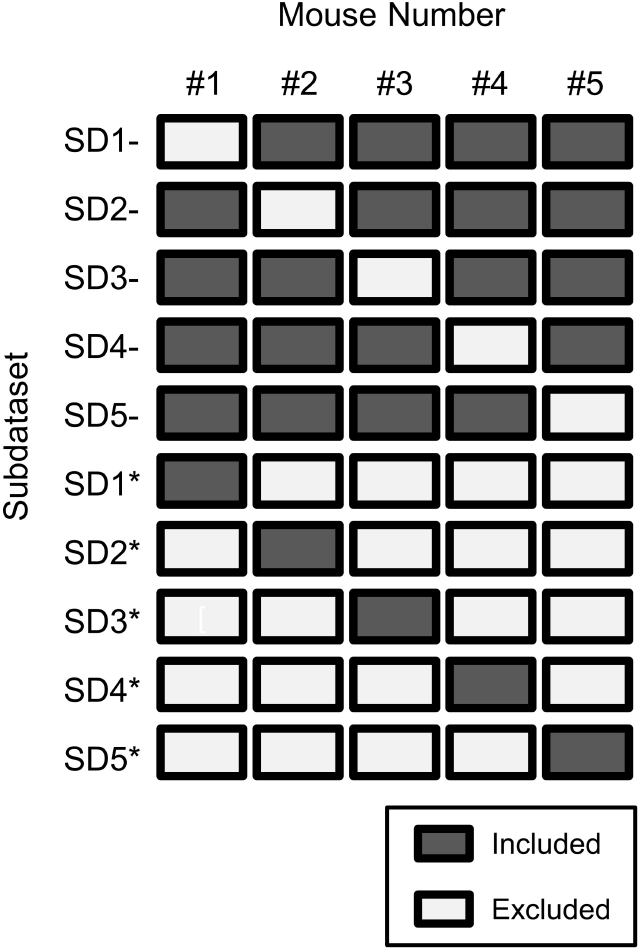
Data splitting for applying Confident Learning. Data were split to define ripples and run Confident Learning and to avoid “data leakage” in the ripple detection experiment. For subdataset SD1-, SD2-, SD3-, SD4-, and SD5-, each mouse data was excluded from the original dataset. For subdataset SD1*, SD2*, SD3*, SD4*, and SD5*, the data obtained for each mouse were treated as a dataset.

Consequently, each ripple candidate was tagged with five kinds of cleaned labels according to the five datasets, including the candidate. Between any pair of datasets, the cleaned labels' concordance rate was ≥ 0.93 (Fig. 9). This result suggested that the cleaned labels were robust for data analysis in Confident Learning, at least based on the applied settings. Additionally, the predicted probabilities for the T_Ripple_CNN_ group for any pair of datasets were positively correlated (Fig. 10; ps < .05, multiple comparisons using the test for no correlation with Bonferroni correction), although there were some comparatively small correlation coefficients (e.g., 0.24).

**Figure 9.**
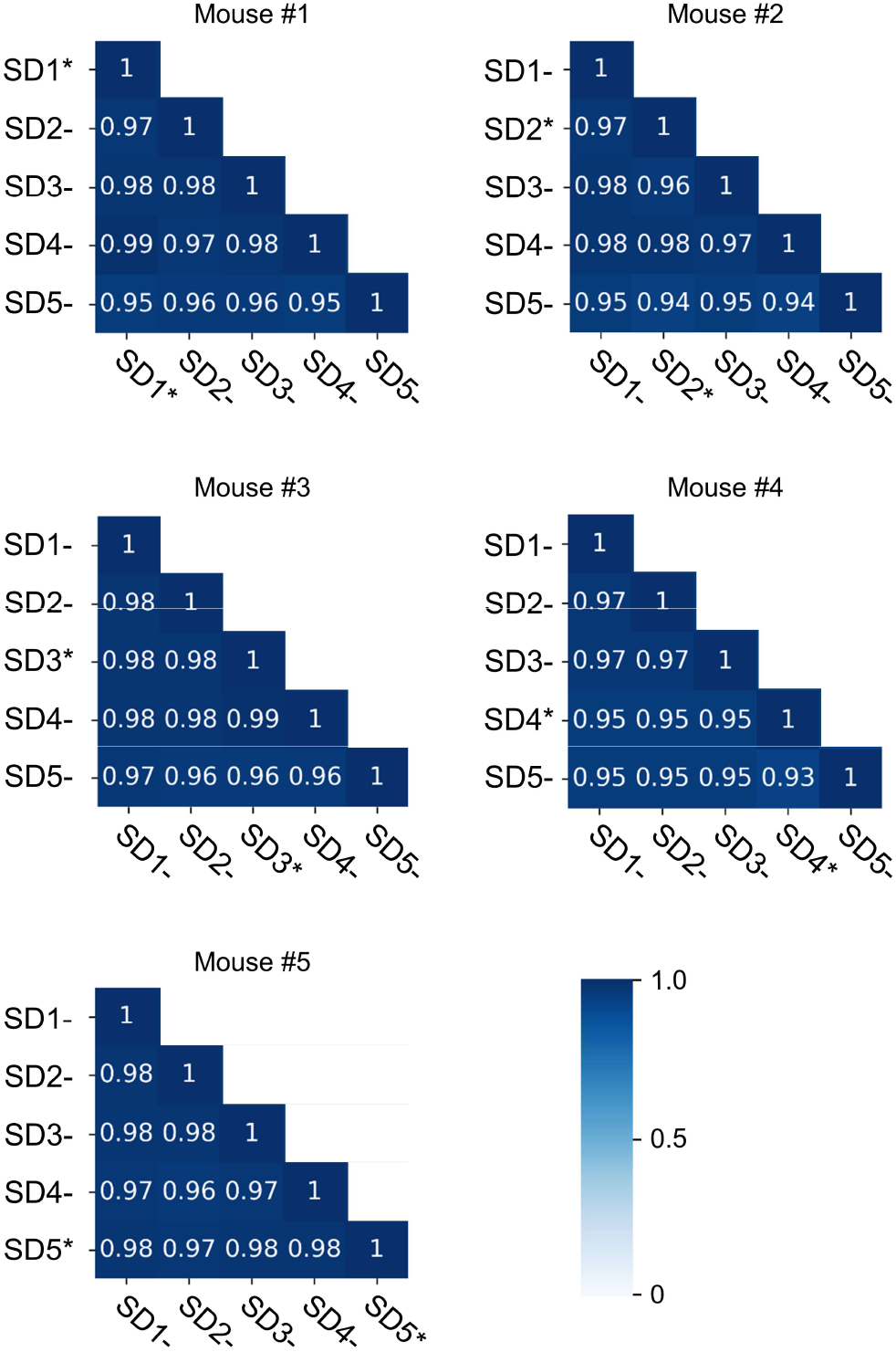
Concordance rates of ripple labels defined with CNNs trained on different datasets. The matrices show the concordance rates of the labels (T_Ripple_CNN_ or F_Ripple_CNN_) estimated with Confident Learning. That is, every ripple candidate was included in five datasets (e.g., a ripple candidate recorded for mouse #1 was included in the subdataset SD1*, SD2-, SD3-, SD4-, and SD5-as shown in Fig. 8). Therefore, each ripple candidate was classified five times as a true-ripple-like event 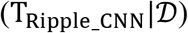 or false-ripple-like event 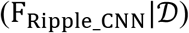 according to the subdatasets.

**Figure 10.**
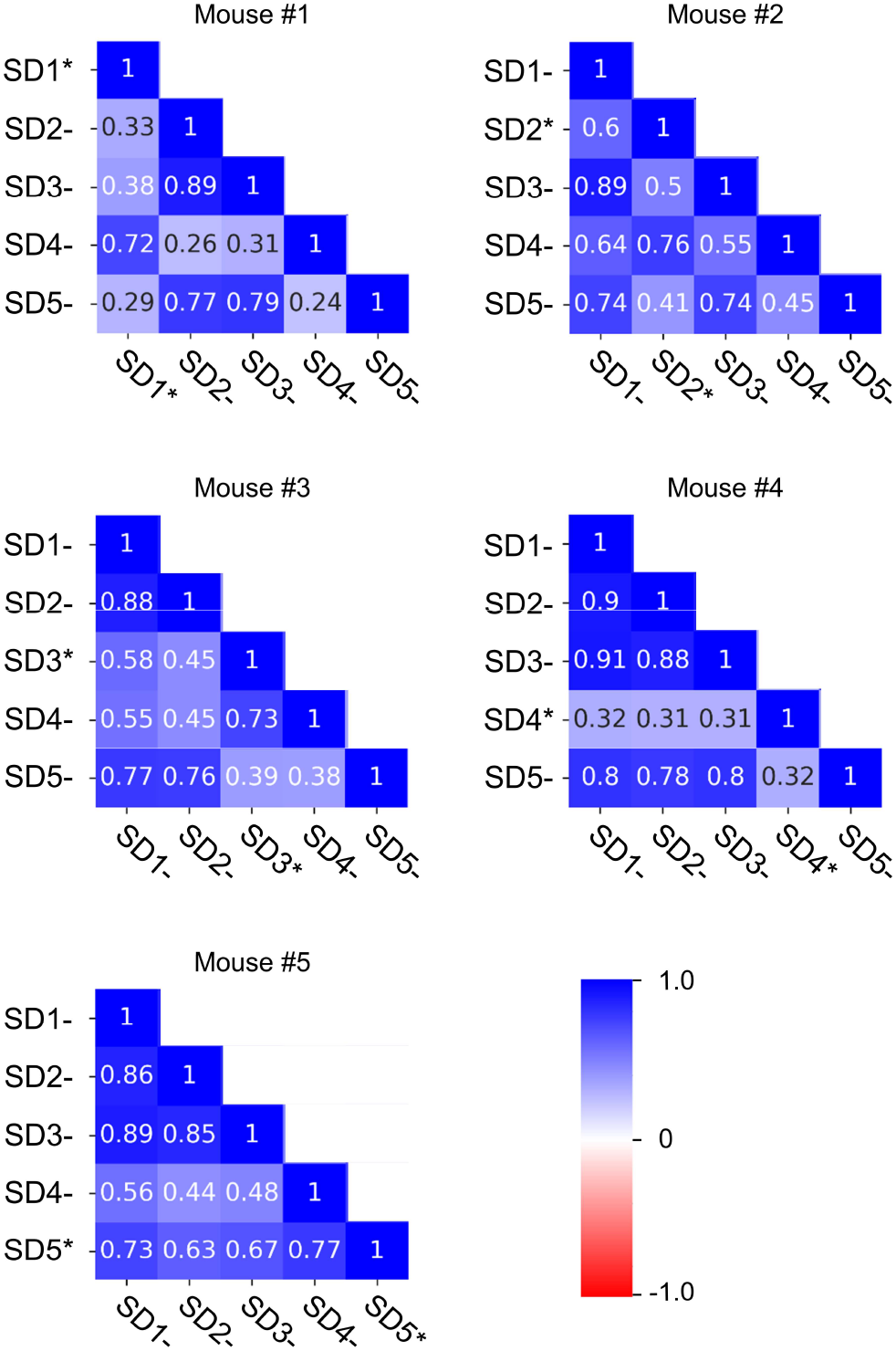
Correlations among estimated probabilities as true ripples based on our CNN and different datasets. The matrices show the Pearson correlation coefficients of predicted probabilities as ripples (= P(**c**_**i**_ ∈ T_Ripple_CNN_) as in Equation (37)) estimated with Confident Learning between subdatasets shown in Fig. 8.

Next, we checked the properties of the ripple candidates subgrouped based on the labels T_Ripple_GMM_, F_Ripple_GMM_, T_Ripple_CNN_ and F_Ripple_CNN_. From the following analyses, we considered cleaned labels from only one dataset. For the ripple candidates defined from the mouse #1, #2, #3, #4, and #5, the cleaned labels determined from using the subdataset SD2-, SD1-, SD1-, SD1-, and SD1-was considered, respectively (Fig. 11). For the noisy and cleaned labels, all four label combinations (2 × 2) were assessed. We named these combinations T2T group (T_Ripple_GMM_ to T_Ripple_CNN_), T2F group (T_Ripple_GMM_ to F_Ripple_CNN_), F2T group (F_Ripple_GMM_ to T_Ripple_CNN_), and F2F group (F_Ripple_GMM_ to F_Ripple_CNN_) (Figs. 12 & 13).

**Figure 11.**
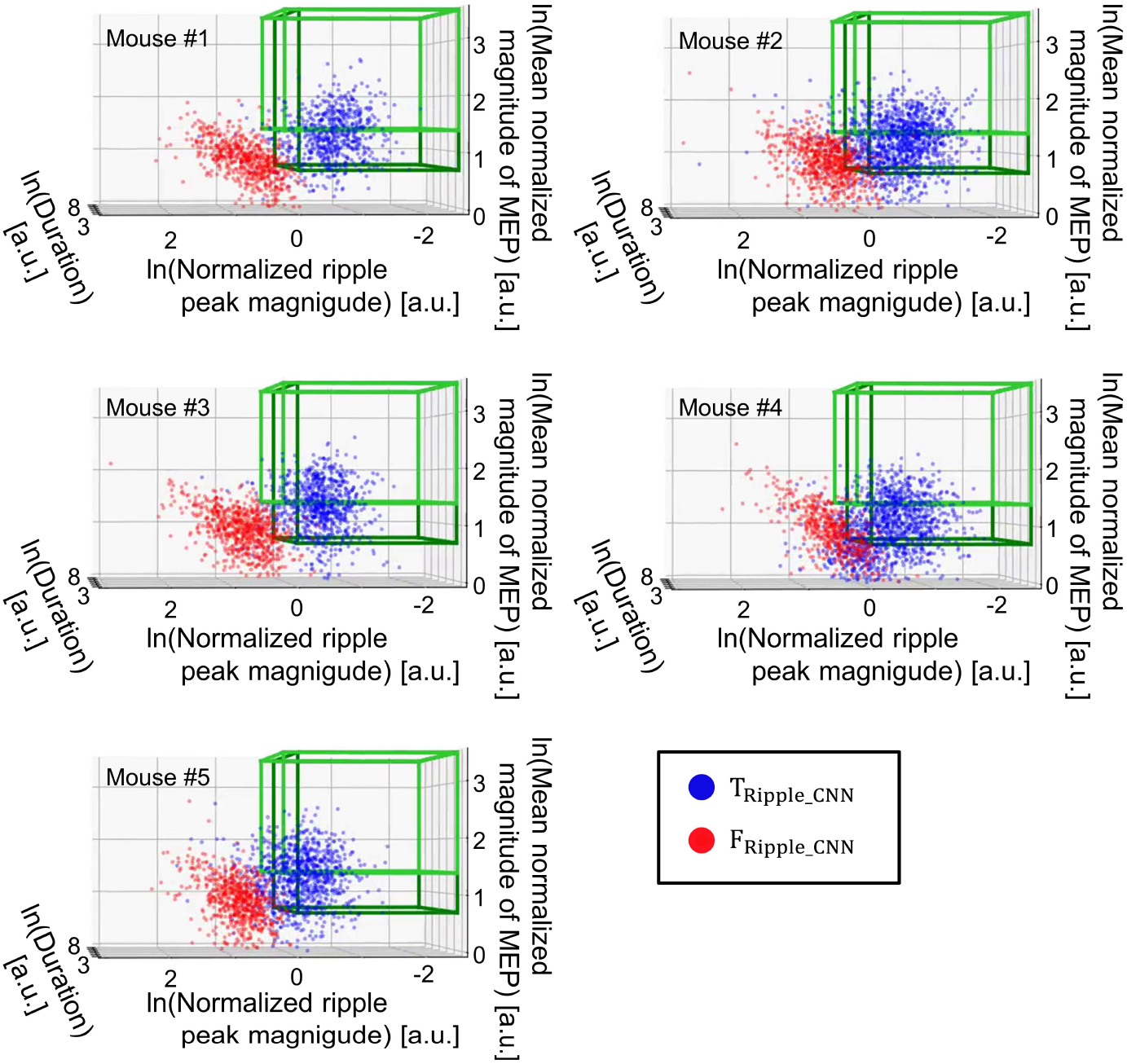
Ripples defined using our CNN. Each plot in the 3D space shows a true ripple-like event (*Blue*; labeled T_Ripple_CNN_) or false ripple-like event (*Red*; labeled F_Ripple_CNN_) defined with our CNN trained in a weakly supervised manner. The data cloud (T_Ripple_CNN_ and F_Ripple_CNN_) and light and dark green cubes are the same as those in Figs. 2 and 5. Note that the input to our CNN was not the three “ln-variables”, as in the case of the Gaussian mixed model, but a 400-ms LFP raw trace. Additionally, the decision boundaries in this 3D space were not hyperplanes, unlike those in Gaussian clustering in Fig. 5.

**Figure 12.**
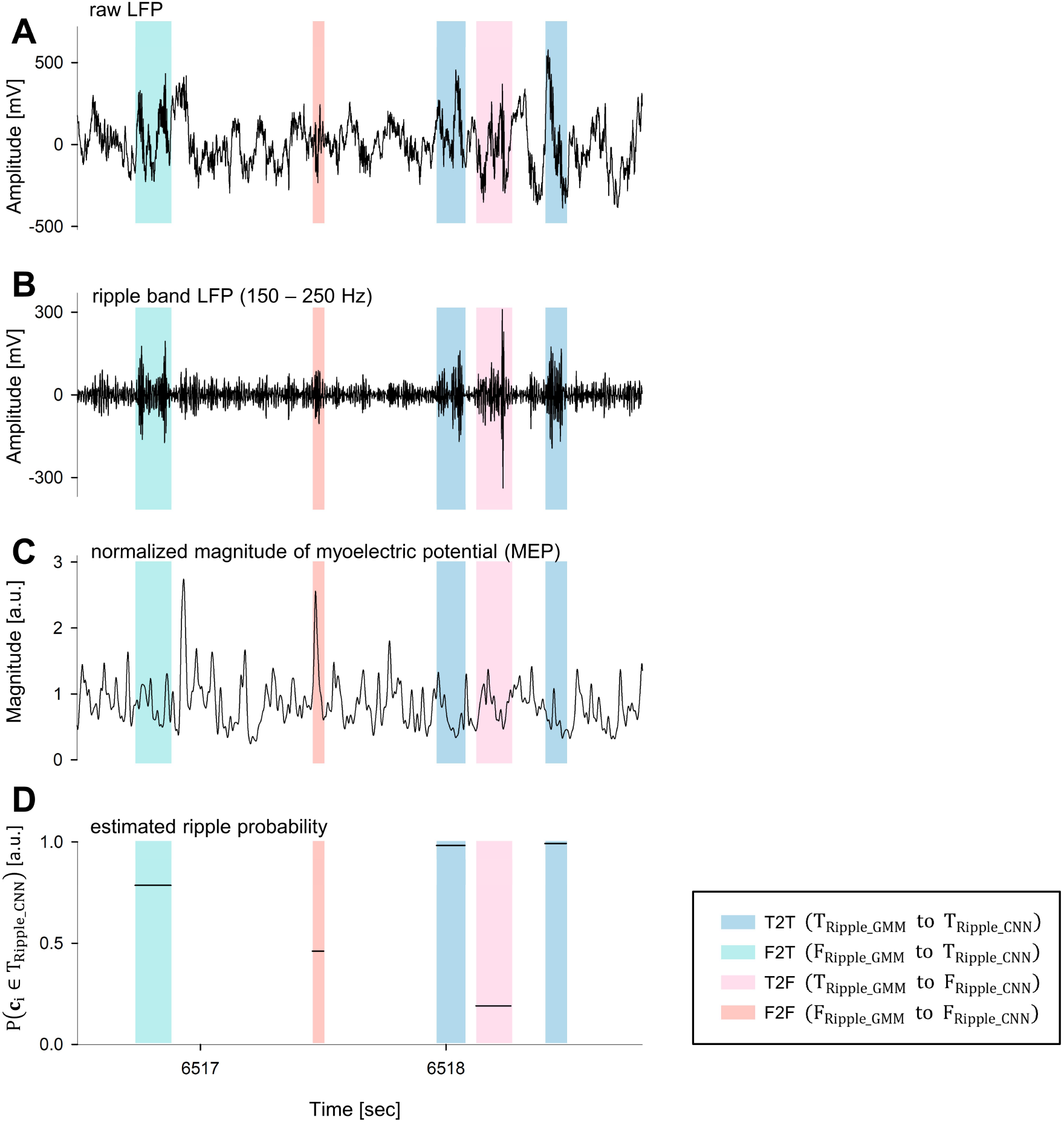
Representative traces of ripple candidates with labels estimated by our CNN. Representative LFP and MEP traces acquired from mouse #1. **A.** Hippocampal raw LFP. **B.** The corresponding trace of the hippocampal ripple band LFP (150–250 Hz). **C.** The corresponding trace of the normalized magnitude of the myoelectric potential (MEP) of the trapezius [a.u.]. **D.** Predicted probabilities estimated by our CNN to define ripples. Note that colored rectangles show the four subgroups of ripple candidates: *Blue*, T2T group (T_Ripple_CNN_ to T_Ripple_CNN_); *Green*, T2F group (T_Ripple_GMM_ to F_Ripple_CNN_); *Pink*, F2T group (F_Ripple_GMM_ to T_Ripple_CNN_); *Red*, F2F group (F_Ripple_GMM_ to F_Ripple_CNN_).

The medians of the “three variables” were different between any two groups for each mouse (Figs. 14A, 15A, and 16A; ps < .01; multiple comparisons using the Brunner-Munzel test with the Benjamini-Hochberg correction after the Kruskal-Wallis test). We also calculated the above comparisons' effect sizes to quantify how substantial the supported differences were, independently of the sample size. Here, we adopted Cliff’s delta statistic (Cliff, 1996) as an index of the effect size because it does not require any assumption regarding the data distribution nor homoscedasticity between two compared groups. Cliff’s delta statistic can range from −1 to 1, and the absolute value indicates the difference; specifically, < 0.147 is negligible, < 0.330 is small, < 0.474 is medium, and ≥ 0.474 is large (Romano, 2006).

**Figure 13.**
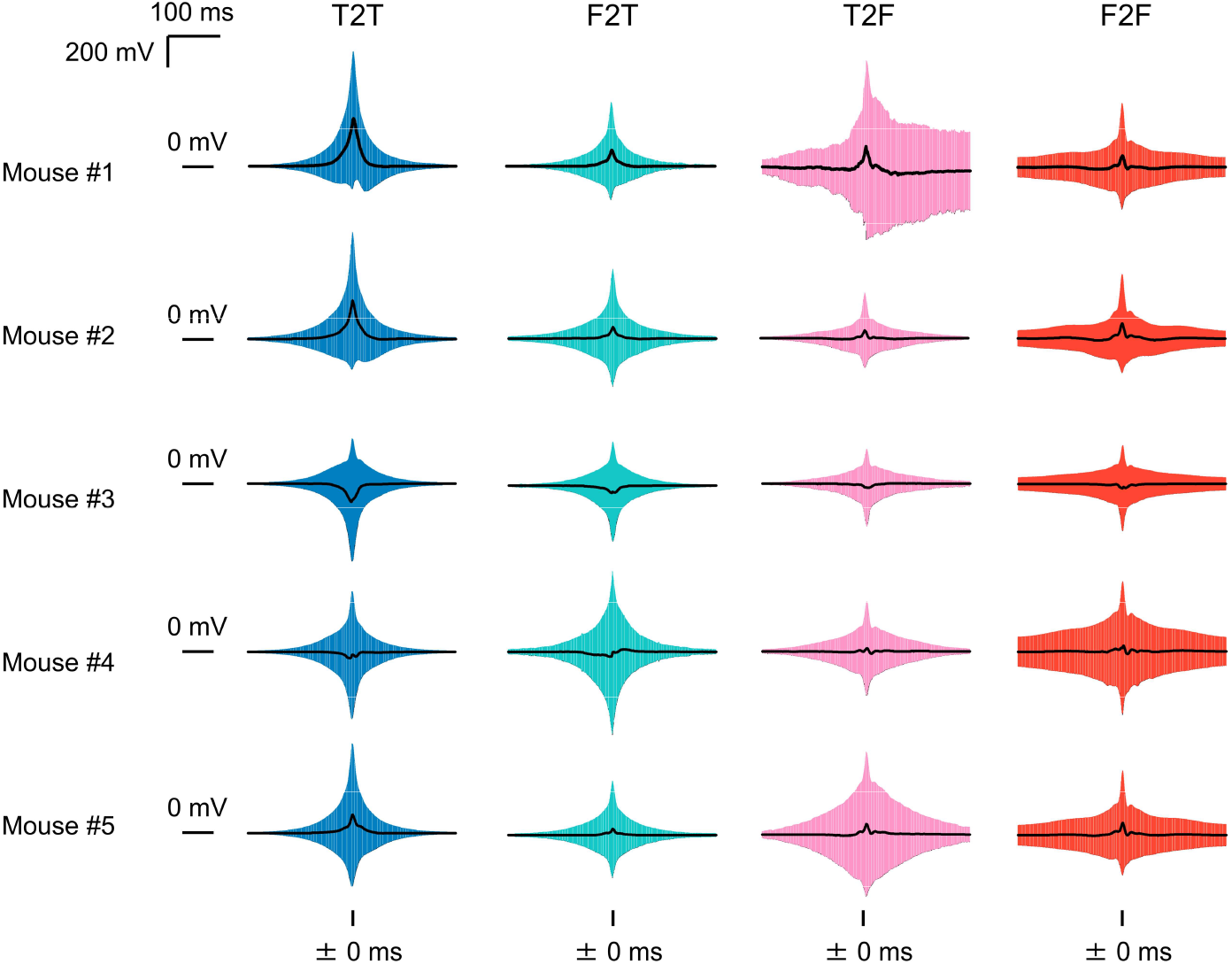
Average traces of ripple candidates. The average traces of the ripple candidates obtained based on the middle time. Rows correspond to each mouse, and columns correspond to the following four subgroups: *Blue*, T2T group (T_Ripple_GMM_ to T_Ripple_CNN_); *Green*, T2F group (T_Ripple_GMM_ to F_Ripple_CNN_); *Pink*, F2T group (F_Ripple_GMM_ to T_Ripple_CNN_); *Red*, F2F group (F_Ripple_GMM_ to F_Ripple_CNN_). Black solid lines show the means. Colored areas express the standard deviations at the corresponding temporal distance from the means.

**Figure 14.**
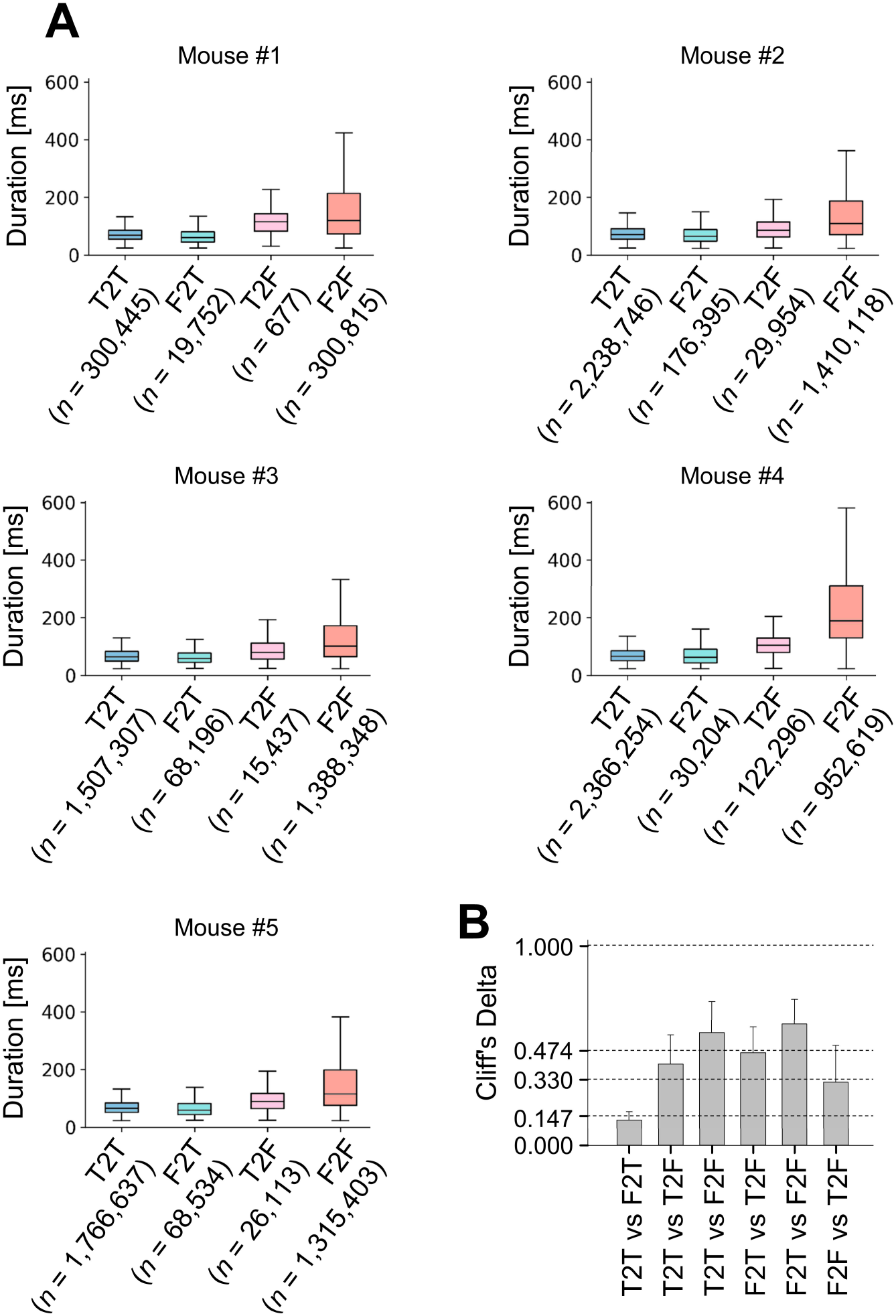
The distributions of the durations among the ripple candidate subgroups. **A.** Boxplots of the duration of each ripple candidate for the ripple subgroups: T2T (T_Ripple_GMM_ to T_Ripple_CNN_; *Blue*), F2T (F_Ripple_GMM_ to T_Ripple_CNN_; *Light Blue*), T2F (T_Ripple_GMM_ to F_Ripple_CNN_; *Pink*), F2F (F_Ripple_GMM_ to F_Ripple_CNN_; *Red*). In all four groups for any mouse, the differences in medians were statistically significant (*ps* < .01; Brunner-Munzel test with the Bonferroni correction after the Kruskal-Wallis test). **B.** Cliff’s delta statistics for the duration of each ripple candidate, as the effect size, between all pairs of the four groups (mean (+/−) std., n = 5 mice).

First, the Cliff’s delta statistic for the durations [ms] of F2T vs T2T, F2F, or T2F groups were 0.13 ± 0.04 (negligible), 0.61 ± 0.12 (large), and 0.47 ± 0.13 (large), respectively (Fig. 14B; n = 5 mice, mean ± std.). These results show that regarding the duration distribution, F2T group, or the ripples found by Confident Learning stage, was not close to F2F nor T2F group but to T2T group. Indeed, the median durations for T2T, F2T, F2F, and T2F group were 66 ± 2, 60 ± 3, 127 ± 31, and 95 ± 13 [ms], respectively (*n* = 5 mice, mean ± std.). Previous research that used multiple electrodes called silicon probes to detect ripples based on oscillation trends showed that the mode of the duration of sharp waves superimposed on ripples was 50 ms in rats (Sullivan et al., 2011, Buzsáki, 2015). Considering the duration profile, the ripples defined based on the proposed method for T2T and F2T group, or T_Ripple_CNN_, were more similar to the ripples defined with the silicon probes in the previous study than were those for the F2F and T2F groups, or F_Ripple_CNN_.

Second, the Cliff’s delta statistic for the mean normalized magnitude of the MEP [a.u.] of F2T vs T2T, F2F, and T2F group were 0.93 ± 0.02 (large), 0.31 ± 0.08 (small), and 0.78 ± 0.08 (large), respectively (Fig. 15B; n = 5 mice, mean ± std.). These results show that regarding the distribution of the mean normalized magnitude of the MEP, F2T group, or the ripples found by Confident Learning stage, was not close to T2T nor T2F group but rather to F2F group, which had the largest median among the four groups. This is different from existing ripple detection methods, which avoids detecting ripples with activity-related noises.

**Figure 15.**
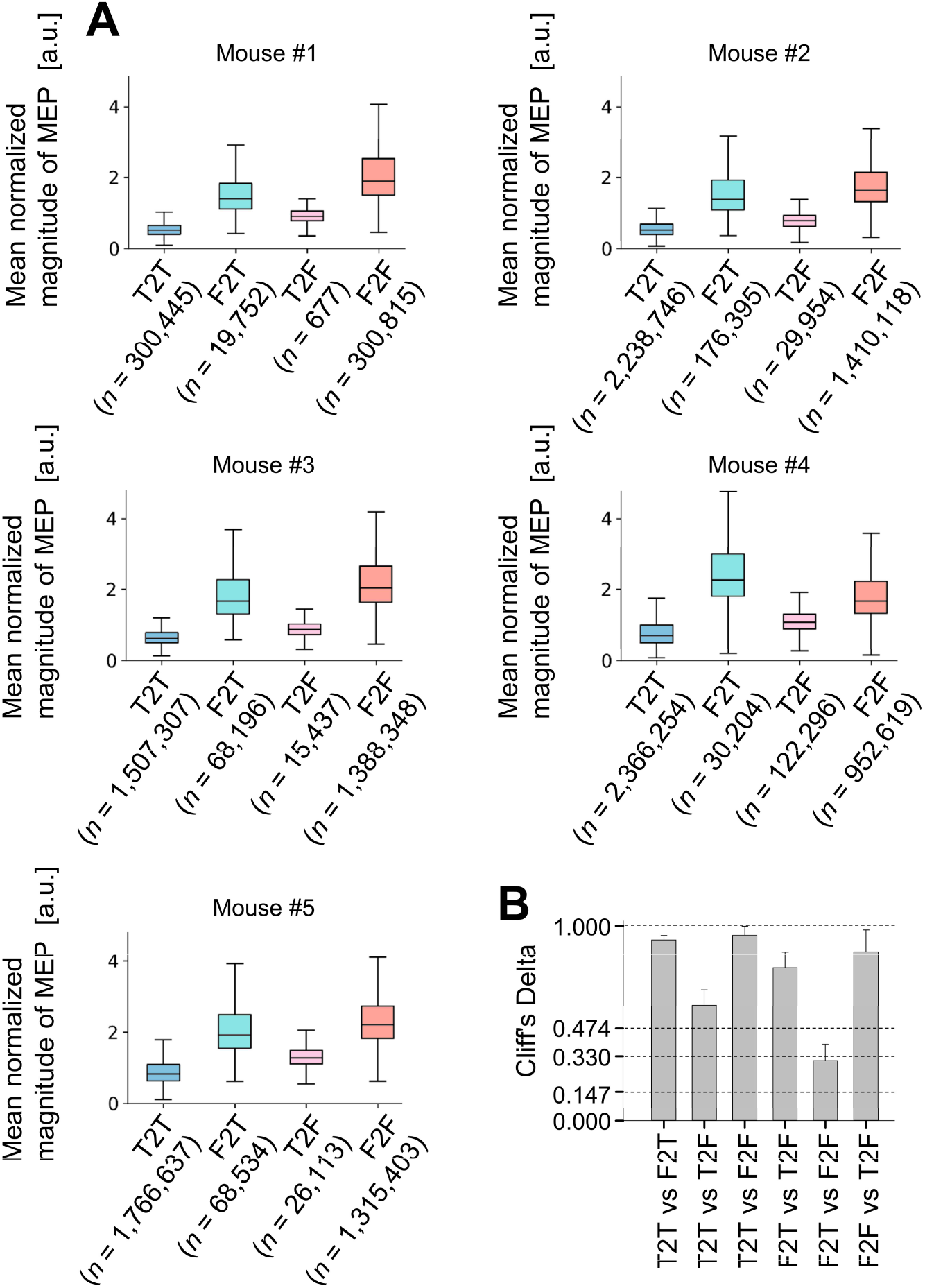
The distributions of the mean normalized magnitude of the myoelectric potential (MEP) among ripple candidate subgroups. **A.** Boxplots of the mean normalized magnitude of the myoelectric potential (MEP) of each ripple candidate concerning ripple subgroups: T2T (T_Ripple_GMM_ to T_Ripple_CNN_; *Blue*), F2T (F_Ripple_GMM_ to T_Ripple_CNN_; *Light Blue*), T2F (T_Ripple_GMM_ to F_Ripple_CNN_; *Pink*), F2F (F_Ripple_GMM_ to F_Ripple_CNN_; *Red*). In all four groups for any mouse, the differences in medians were statistically significant (ps < .01; Brunner-Munzel test with the Bonferroni correction after the Kruskal-Wallis test). **B.** Cliff’s delta statistics for the mean normalized magnitude of the myoelectric potential (MEP) of each ripple candidate, as the effect size, between all pairs of the four groups (mean (+/-) std., n = 5 mice).

Third, the Cliff’s delta statistic for the normalized peak magnitude [a.u.] of F2T vs F2F groups was 0.13 ± 0.08 (negligible) (Fig. 16B; n = 5 mice, mean ± std.). This result indicates that ripple peak magnitude was not a dominant factor of the classification on Confident Learning stage.

**Figure 16.**
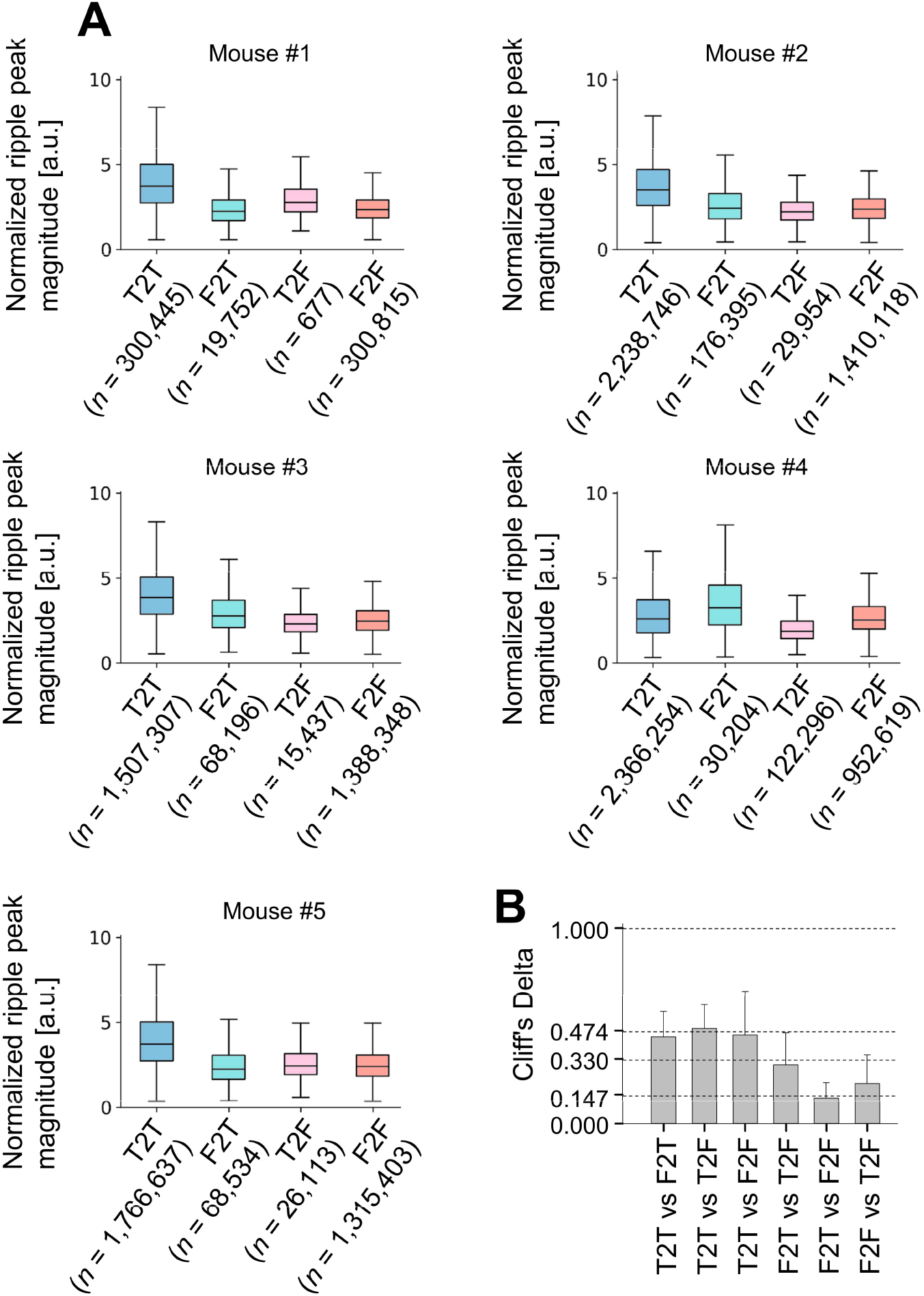
The distributions of the normalized ripple peak magnitudes among ripple candidate subgroups. **A.** Boxplots of the normalized ripple peak magnitudes for each ripple candidate concerning ripple subgroups: T2T (T_Ripple_GMM_ to T_Ripple_CNN_; *Blue*), F2T (F_Ripple_GMM_ to T_Ripple_CNN_; *Light Blue*), T2F (T_Ripple_GMM_ to F_Ripple_CNN_; *Pink*), F2F (F_Ripple_GMM_ to F_Ripple_CNN_; *Red*). In all four groups for any mouse, the differences in medians were statistically significant (ps < .01; Brunner-Munzel test with the Bonferroni correction after the Kruskal-Wallis test). **B.** Cliff’s delta statistics for the normalized ripple peak magnitude for each ripple candidate, as the effect size, between all pairs of the four groups (mean (+/−) std., n = 5 mice).

In summary, the F2T group reflects the phenomenon of hippocampal ripples, and the proposed method succeeded in defining ripples even during movement at a stochastic scale. Notably, the distribution of the mean normalized magnitude of the MEP for F2T group was substantially shifted upward compared with that of the T2T group; that is, Cliff’s delta between the two groups was 0.93 ± 0.02 (Fig. 15B; *n* = 5 mice, mean ± std.).

### Detecting Ripples by Using Our CNN

Next, we determined whether our CNN detects “ripples” from unseen LFP signals (here, “ripple” is a ripple defined using our method). Specifically, we determined whether our CNN could classify 400-ms raw LFP signals into two classes: a ripple-not-including group and a one-ripple-including group, which were labeled R_0_ and R_1_, respectively (Fig. 17A). To evaluate the results, we used the leave-one-animal-out cross-validation method (Fig. 17B).

**Figure 17.**
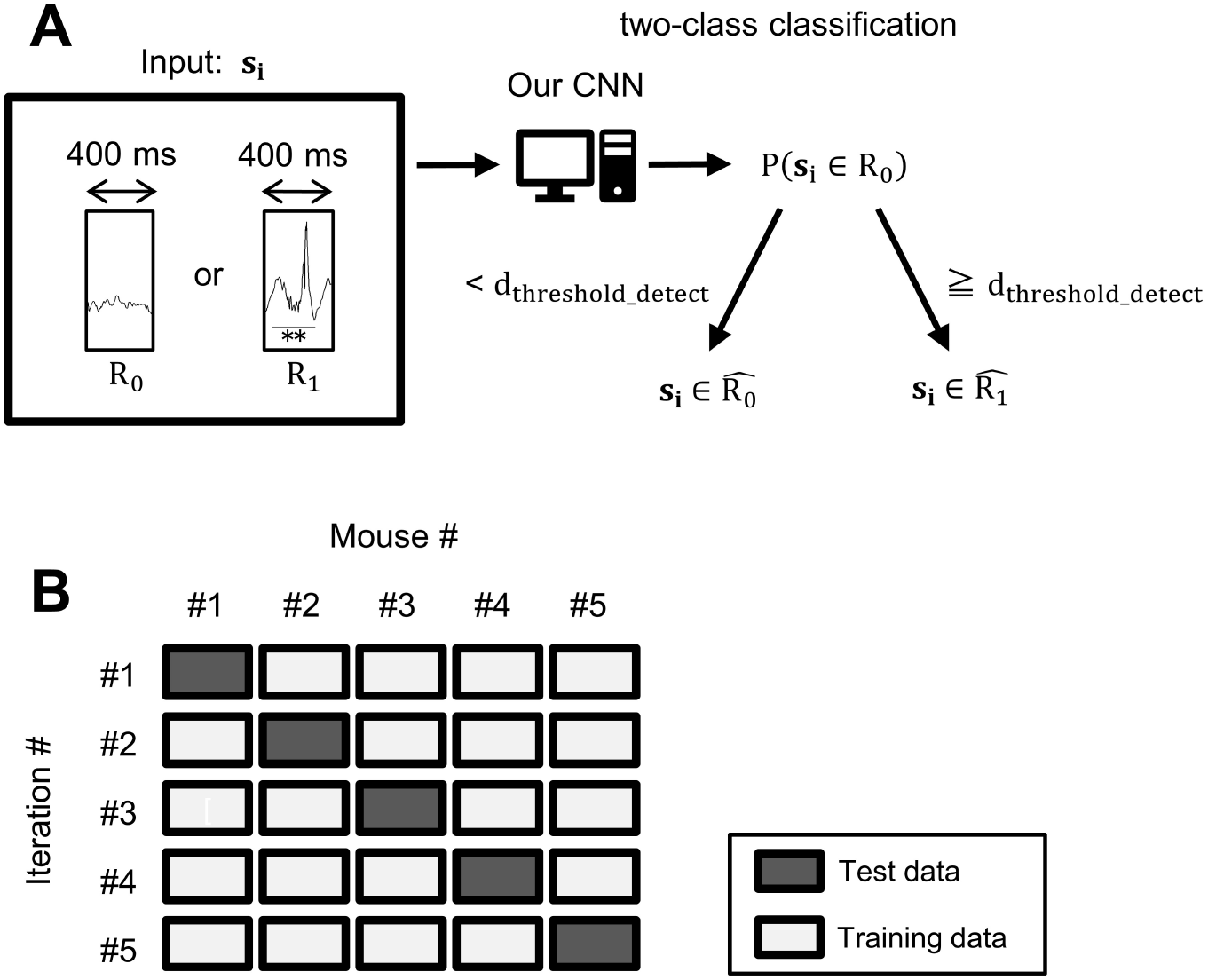
Experimental scheme for detecting ripples. **A.** The experimental design for ripple detection using our CNN. Our CNN requires the input signal vector (**S**_i_) to be a fixed length, 400 points (e.g., 400 ms for a 1 kHz sampling rate) in this study. Each input signal **S**_i_ was separated into the ripple-not-including group (R_0_) or the one-ripple-including group (R_1_). The ripple-not-including group (R_0_) consisted of the 400-ms raw LFP signals that did not include any parts of the ripple candidates. The one-ripple-including group (R_1_) consisted of the 400-ms raw LFP signals, each of which included just one “reasonable ripple,” which was tagged as T_Ripple_CNN_ and had a ripple peak magnitude that exceeded 7 SD. The output of our CNN, P(**S**_i_ ∈ R_1_), was interpreted as the probability that the input to belonged to the one-ripple-including group (R_1_), which was binarized by a decision threshold (d_threshold_detect_) when necessary. **B.** Data splitting was performed based on the leave-one-animal-out cross-validation method. In each iteration, data obtained for each mouse were used as test data.

First, we defined two groups: the ripple-not-including group (R_0_) and one-ripple-including group (R_1_). The ripple-not-including group (R_0_) included 400-ms raw LFP signals that did not include any part of ripple candidates. The one-ripple-including group (R_1_) included 400-ms raw LFP signals, each of which included just one “reasonable ripple,” which was tagged as T_Ripple_CNN_ and had a ripple peak magnitude that exceeded 7 SD. Note that for simplicity, we excluded 400-ms raw LFP signals that included any part of more than two ripples from this experiment. The threshold of the ripple peak magnitude, 7 SD, was determined based on the observation from Fig. 16 that the ripples with ripple peak magnitudes above this level were especially reliable; notably, supervised learning should be performed with highly dependable labels.

As a result, the area under the precision-recall curve for detecting a “reasonable ripple” in each 400-ms raw LFP signal was 0.72 ± 0.10 (Fig. 18; n = 5 mice, mean ± std.). Since this score exceeded the chance level of 0.50, our trained CNN was validated. Here, the output of our CNN was a stochastic scale, and a decision threshold was used to obtain predicted labels (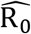 or 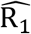; Fig. 17A). For example, if setting the decision threshold to 0.5 (since the task was a two-class classification problem), predicted labels were determined (as Equation (38)), and confusion matrices were obtained (Fig. 19). Regarding the one-ripple-including group (R_1_), the precision, recall, and F1-score were 0.15 ± 0.13, 0.96 ± 0.03, and 0.24 ± 0.18, respectively (Fig. 20; n = 5 mice, mean ± std.). Although the sample size of the one-ripple-including group (R_7_) was less than 1/100 of that of the ripple-not-including group (R_0_), the precision for detecting a ripple was smaller than the recall. This result was caused by our original cost-sensitive learning design for imbalanced data (see Materials and Methods, Equations (42)–(47)). In fact, without the modification of the original cross-entropy loss function, the predictions of our CNN for ripple detection all fell within the majority group, i.e., the ripple-not-including group 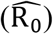.

**Figure 18.**
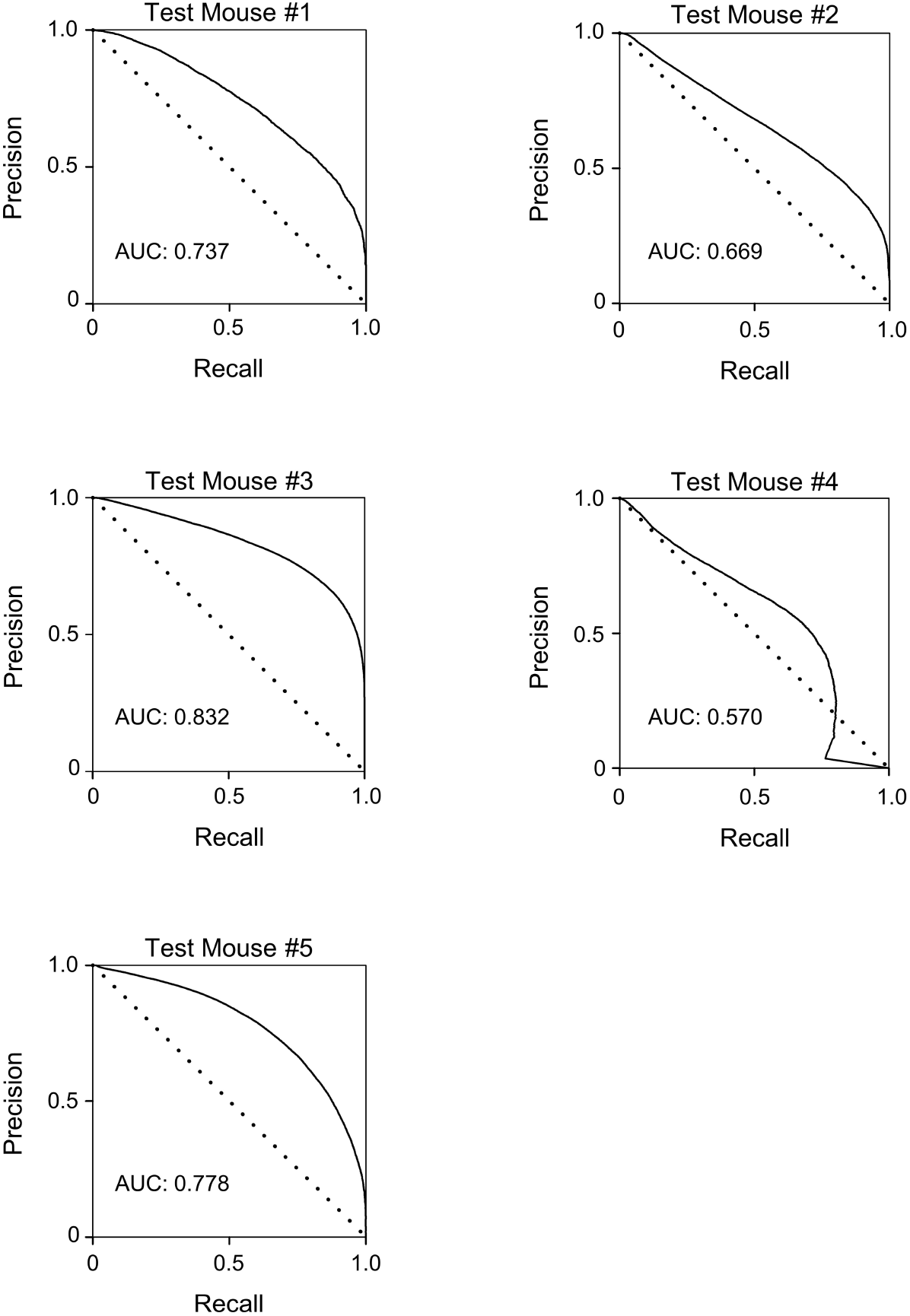
Precision-recall curves for ripple detection. Each panel shows the precision-recall curve for each fold in the leave-one-animal-out cross-validation method. The curves were calculated from the precision and recall results for ripple detection based on when the decision threshold changed from zero to one (see Fig. 17 and Equation (41) in Materials and Methods). The AUC (area under curve) ranged between zero and one and reflects the two-class classification ability of the proposed approach averaged over the decision thresholds; this approach is appropriate for imbalanced datasets, such as in this case.

**Figure 19.**
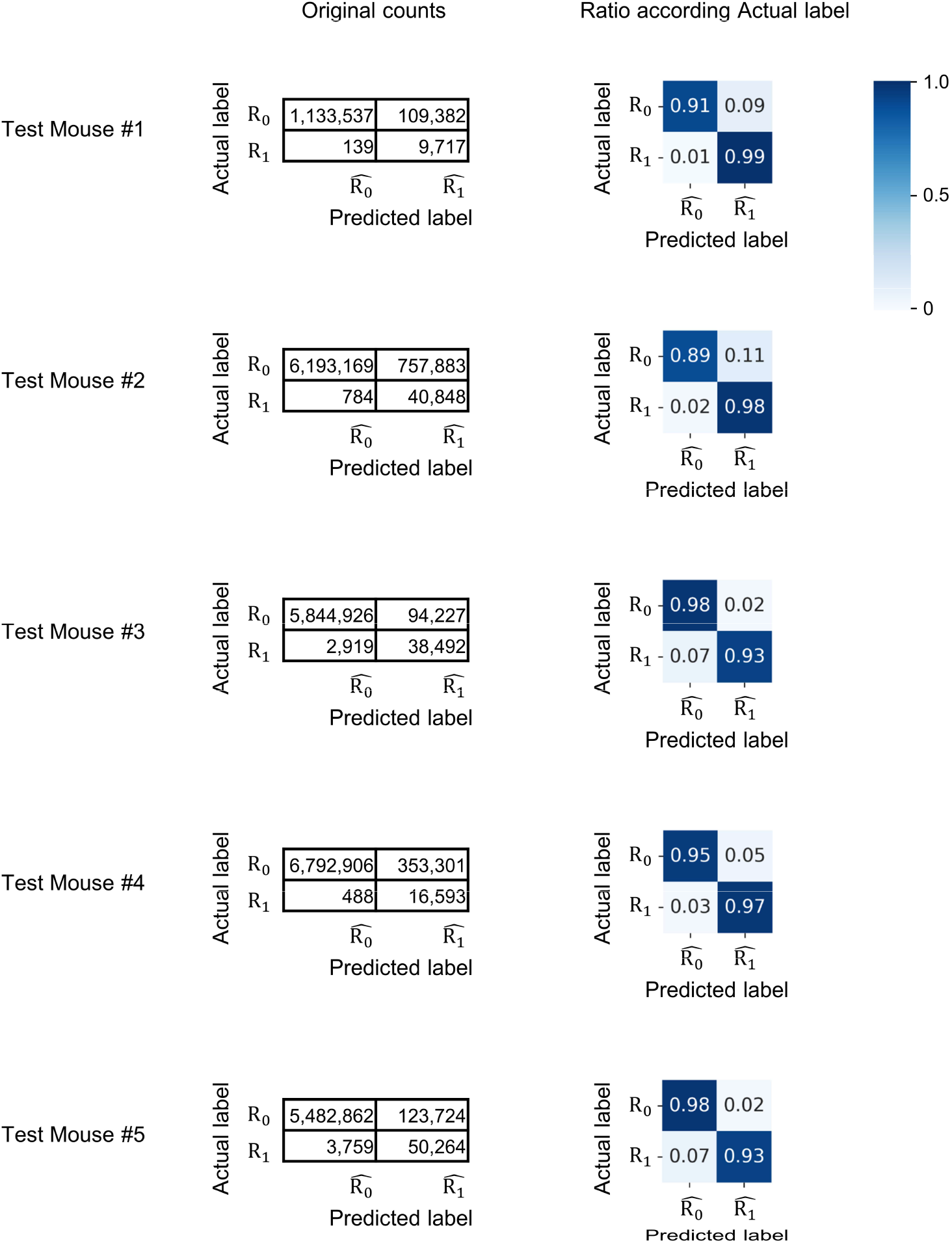
Confusion matrices for ripple detection. The panels show confusion matrices for ripple detection when the binarization threshold is fixed to 0.5 (see Fig. 17A and Equation (41) in Materials and Methods). Each 400-ms raw LFP sample was classify into two groups: a ripple-not-including group (R_0_) and a one-ripple-including group (R_1_). Each row shows the confusion matrices for each fold in the leave-one-animal-out cross-validation method (*Left Column*: original values, *Right Column*: values normalized by the sample sizes of actual groups).

**Figure 20.**
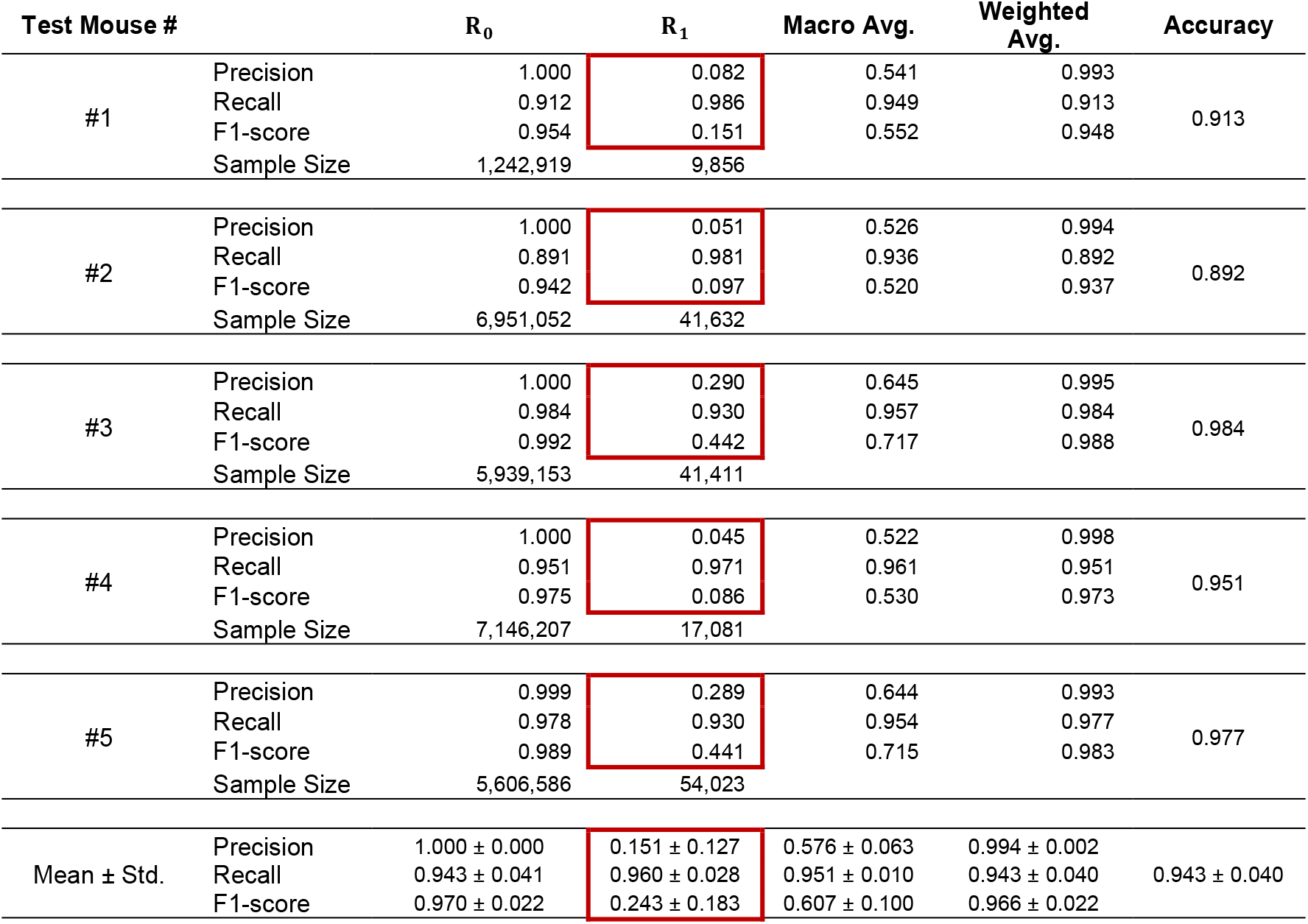
Metrics for detecting ripples. A summary of the metrics for detecting ripples when the decision threshold was fixed to 0.5 is given. Note that the F1-score of ripple detection (surrounded by the red rectangle) can be optimized by searching for optimal decision thresholds (as Equation (41)) using validation data. However, we did not try that because of the difficulty of splitting data from five mice into training, validation, and test datasets.

### Reverse Estimation of the Optimal Threshold Based on the Ripple Peak Magnitude

There has been considerable debate regarding the best threshold value for the ripple peak magnitude in existing methods of ripple detection. Here, we explored whether there is an optimal threshold that would maximize the F1-score for detecting ripples using the trained and validated CNN obtained from the previous experiment.

To search for the optimal threshold, we used “suspicious ripples,” which were defined as ripples labeled as T_Ripple_CNN_ with ripple peak magnitudes from 1 SD to 7 SD. Additionally, we defined one-suspicious-ripple-including group (R_s_) that included 400-ms raw LFP signals, each of which included just one “suspicious ripple.” The optimal threshold of the ripple peak magnitude was determined by using the trained CNN to separate the one-suspicious-ripple-including group (R_s_) into the ripple-not-including group (R_0_) and the one-ripple-including group (R_1_).

First, we plotted the probabilities predicted by the trained CNN for samples from the one-suspicious-ripple-including group (R_s_) that belonged to the one-ripple-including group (R_1_) (Fig. 21). From 1 to 10 SD, the range of the ripple peak magnitude [mV] was split into 0.1 SD bins without overlaps. For each bin, the mean and standard deviation of the probabilities predicted by the trained CNN were calculated.

**Figure 21.**
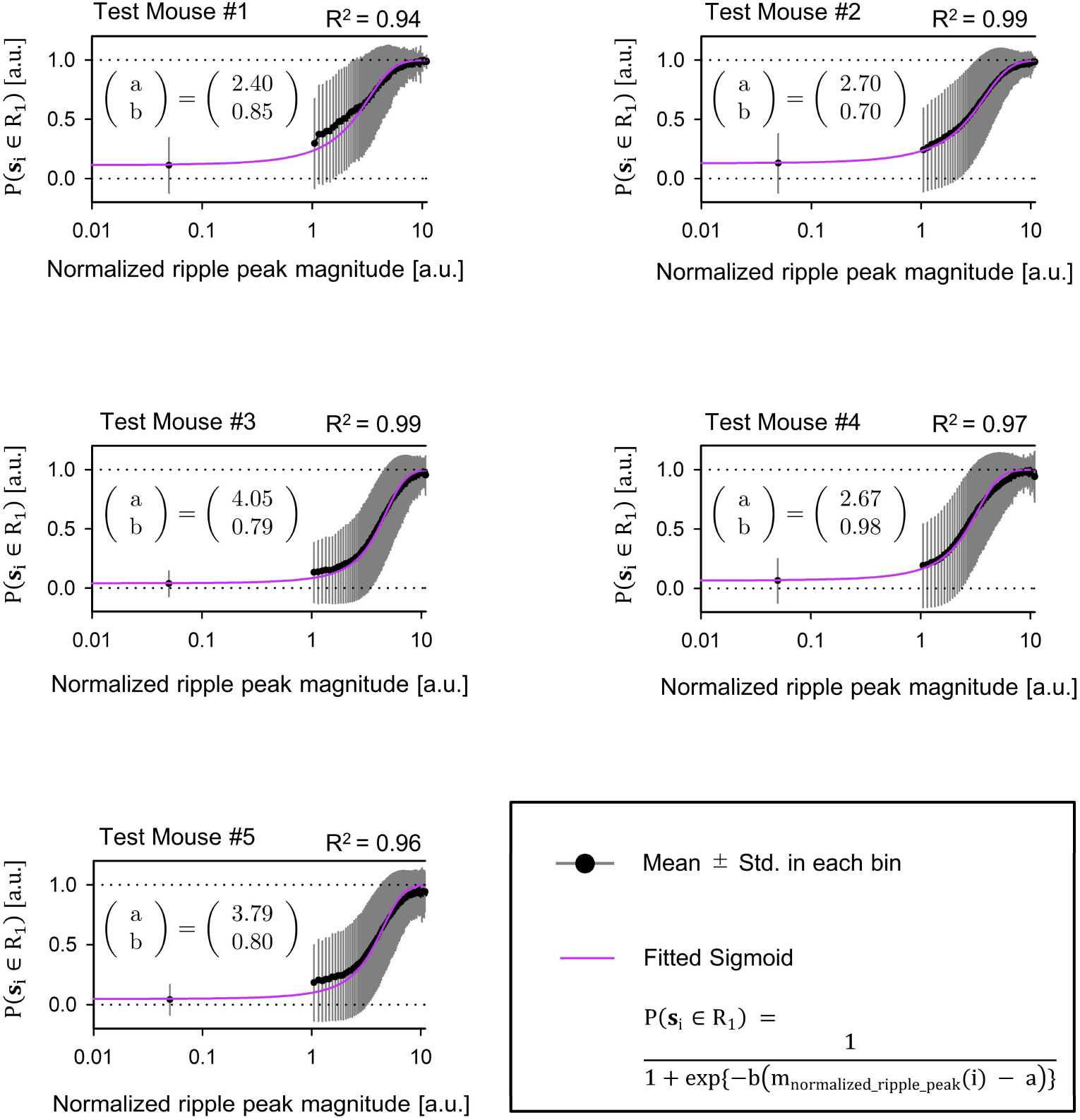
Reverse estimation of the optimal threshold based on the ripple peak magnitude. The optimal threshold for ripple detection based on the ripple peak magnitude was reversely estimated using our CNN. This approach was verified to effectively detect ripples at AUCs higher than those attainable by chance, as shown in Fig. 18. From 1 SD to 10 SD, the range of the ripple peak magnitude [mV] was split into bins every 0.1 SD without overlaps. We trained the CNN to predict samples from the one-suspicious-ripple-including group (R_s_) to belong to the ripple-not-including group (R_0_) or one-ripple-including group (R_1_). Sigmoid functions (*Purple*, see Equation (49) in Materials and Methods) were fit based on the mean predicted probabilities for the one-ripple-including group (R_1_) for each bin (the coefficient of determination R^2^ = 0.97 ± 0.02, mean ± std., *n* = 5 mice).

Moreover, a sigmoid function, including parameters a and b, was fit based on all observed data pairs.

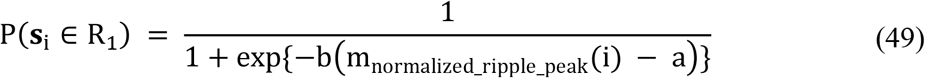

where m_normalized_ripple_peak_(i) is the normalized ripple peak magnitude [a.u.] of the i-th suspicious ripple based on Equations (1)–(5) and (11)–(14), ***s***_**i**_ is the 400-ms raw LFP signal that includes the i-th suspicious ripple, and a and b are the parameters used for fitting. Specifically, a is the normalized ripple peak magnitude value predicted by the fitted sigmoid function at the inflection point, and b is the slope of the fitted sigmoid function at the inflection point as

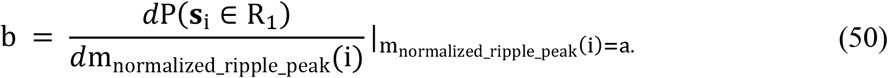

The fitted values a and b were determined to be 3.12 ± 0.74 and 0.83 ± 0.10, respectively (n = 5 mice, mean ± std.). Additionally, the mean predicted probabilities in every bin and the determined sigmoid function yielded the coefficient of determinations R^2^ of 0.97 ± 0.02 (n = 5 mice, mean ± std.), indicating reasonable agreement. These results suggested that setting the ripple peak magnitude to 3.12 ± 0.74 maximizes the F1-score for detecting ripples based on existing ripple detection methods.

## Discussion

Our proposed method defined ripples using a CNN trained in a weakly supervised manner based on “noisy labels” obtained through GMM clustering. These two machine learning steps made it possible to utilize the temporal local features of LFPs and ripples. In the detection stage, our CNN's input was a series of 400-ms hippocampal raw LFPs recorded with one electrode. The output was the probability of the input including a “true” ripple.

From the perspective of duration, the ripples defined by our method (T_Ripple_CNN_ or T2T and F2T group) were closer to the ripples defined by a multielectrode method in a previous study than were the false ripples defined by our method (F_Ripple_CNN_; T2F and F2F group). Additionally, concerning the mean normalized magnitude of the MEP of the trapezius [a.u.], F2T group was larger than that of T2T group. Thus, with Confident Learning, the proposed method enables the detection of ripples during both stable and moving states, the latter of which is not possible with existing methods. Previous studies reported that ripples occur even when animals feed or walk (Buzsáki, 2015). Our approach offers a way to define/detect ripples during active states with just a single electrode.

Since the proposed method defines ripples probabilistically, it is not needed to sort out small ripple candidates. With our approach, binarized predicted labels (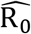: ripple-not-including, or 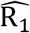: one-ripple-including) can be obtained by adjusting the cut-off parameter of the ripple probability, without losing small ripples using a predefined threshold for the ripple band peak magnitude.

Furthermore, the capacity for handling small ripples contributes to the enhanced robustness of detecting ripples at the recording site in the hippocampus and extending the time limitation for continuously and stably detecting ripples. The LFP amplitude is inversely proportional to the distance between the virtual origin and the recording sites (Buzsáki et al., 2012). Also, it is known that recording electrodes tend to “slip,” especially in the electrode's longitudinal direction under freely moving conditions, at least because brain tissue is characterized by elasticity. Under these settings, our method can be effectively applied to detect ripples, especially in experiments at day-to-week scales. The quality enhancement of ripple detection in terms of stabilization and prolongation is ideal for revealing the relationships between hippocampal ripples and memory consolidation or long-term memory in detail.

From the viewpoint of conventional digital signal processing (DSP) or time-frequency analysis, the length of the convolution filters (K), a hyperparameter of our CNN, is related to the frequency bands' limits to extract features based on convolution.

Our CNN extracted temporal local features over 37 consecutive ms periods. In our settings, 7-, 5-, and 3-unit length convolution filters were included in each of three blocks of our CNN, and the effective sampling rate of the hippocampal LFP was 1 kHz. Therefore, 37 ms (= 1 + {(7 - 1) + (5 - 1) + (3 - 1)} × 3 ms) was the total consecutive time in which local features were extracted with our CNN.

It is possible that low-frequency components, which are difficult to express in 37 ms, may not be treated by our CNN. In fact, ripples (150–250 Hz) often overlap with sharp waves (5–50 Hz; O’Keefe, 1976; Buzsaki, 2015; Watanabe, 2017) and gamma waves (25–75 Hz; Ramirez-Villegas et al., 2015).

If these low-frequency components are not utilized, the following three modifications should be made: (i) enlarge each filter length and (ii) use the Inception module (Szegedy et al., 2014), or (iii) like the SincNet (Ravanelli et al., 2019) to add a relatively long bandpass filter to the first layer of the neural network.

Our method has a limitation; for simplicity, it is assumed that there is not more than one ripple event within a 400-ms LFP input sequence for our CNN in the detection step. However, this limitation is not practical. Ripples often appear in clusters (i.e., events with < 100 ms intervals between them). The proportion of clustered ripples is approximately 50% during brief pauses and approximately 20% during prolonged periods of immobility while awake or periods of sleep in home cages (Buzsáki, 2015).

Our CNN to detect ripples should be modified to detect when and how many ripples in a time series. This refinement might be realized by mimicking object detection models in 2D or 3D. Specifically, the faster RCNN model (Ren et al., 2015) or the CenterNet model (Zhou et al., 2019) may be potential candidates. When using the 1D modified versions of these models, it is needed to balance the relationship among the following three components: the convolutional filter's length, the maximum frequency band to extract features, and the temporal resolution (because of the Gabor uncertainty).

## Acknowledgments

This work was supported by JST ERATO (JPMJER1801) and JSPS Grants-in-Aid for Scientific Research (18H05525). Y.W. and Y.I. conceptualized the study; M.O. performed the experiments; Y.W. performed the data analysis; Y.W. wrote the original draft; and all authors reviewed and edited the final manuscript.

